# Primary and Supplementary Motor cortex implement parallel control solutions for rhythmic and discrete arm movements

**DOI:** 10.64898/2026.02.16.706176

**Authors:** Andrea Colins Rodriguez, Rómulo Fuentes-Flores, Mark Humphries

## Abstract

Arm movements are rhythmic, discrete, or some combination of the two. Conflicting evidence supports each of two possible solutions for how motor cortex controls them: that either it uses the same strategy for controlling rhythmic and discrete movements or different strategies for distinct movement types. Using recurrent neural network modelling and multi-unit recordings during an arm-cycling task, we show that primate motor cortex uses both solutions. Primary motor cortex (M1) dynamics converge to the same limit-cycle when executing both movement types. In contrast, supplementary motor area (SMA) dynamics diverge according to the type of the upcoming movement before reaching a helical spiral. Our results reconcile opposing views on the cortical control of rhythmic and discrete movements by showing that the two solutions are not mutually exclusive but implemented in parallel within motor cortex. Instead, our results challenge us to understand why both solutions are implemented to solve the same problem.

## 1 Introduction

We can generate a myriad of arm movements, from stereotypical rhythmic movements – like whipping cream – to completely discrete movements, like throwing a dart. Despite our ability to generate arm movements that span the rhythmic-to-discrete spectrum [1, 2], how motor cortex activity generates that spectrum is unclear. The few studies that have characterised cortical activity during both types of arm movements in the same task have identified that motor cortex is engaged in both types [3–7], but whether it uses the same or different control strategies to generate discrete and rhythmic arm movements remains unresolved.

There are two possible, contrasting hypotheses for the cortex’s solution to controlling the rhythmic-to-discrete spectrum. On the one hand, discrete and rhythmic arm movements could share the same control strategy. fMRI recordings of motor cortex during wrist movements in humans show that both M1 and SMA are active during the production of both movement types, albeit to a different extent [3]. At the neuronal level, the same-control hypothesis is supported by the similarities between the activity of individual neurons in the primary motor cortex (M1) across both types of movements [5]. At the neural population level, M1 shares dynamical motifs during the production of discrete and rhythmic arm movements, such as rotational dynamics [8–10]. On the other hand, motor cortex could have different strategies for controlling discrete and rhythmic movements. The different-control hypothesis is supported by the differences in the population activity of the supplementary motor area (SMA) across the two types of movement, with helical dynamics during rhythmic arm movements [9] and non-rotational dynamics during discrete movements [11]. Recent recurrent neural network (RNN) models have assumed different control strategies by training the population dynamics controlling discrete and rhythmic movements to occupy independent neural subspaces [12, 13].

To test whether motor cortex employs the same or different strategies to control rhythmic and discrete movements, we explored the neural activity of M1 and SMA while non-human primates performed arm movements that spanned the rhythmic-to-discrete spectrum. We used RNNs to predict the population dynamics of both regions during an arm-pedalling task [9] under each hypothesis for the control of arm movement: either the same control strategy for all movement types (same-control RNNs) or different control strategies for rhythmic and discrete movements (different-control RNNs). We defined the key difference between same- and different-control strategies by the contextual information that the RNNs receive. While same-control strategies consider the number of cycles to be a continuum, the different-control strategies do not. Rather, they receive explicit contextual information defining whether the type of movement is discrete or rhythmic. We found distinct predictions for M1 and SMA, which we tested against neural recordings. The population dynamics of M1 were consistent with a single control strategy that generates both types of movements, with the population dynamics for discrete movements converging to the same limit cycle that generates rhythmic movements. In contrast, SMA’s population dynamics were consistent with it using different control strategies for discrete and rhythmic movements, as population dynamics differed by the type of movement performed, even before its onset. In this way, our results reconcile opposing views of the control of rhythmic and discrete movements by showing that two motor cortical regions employ the same and different control solutions in parallel to address the same problem.

## 2 Results

To characterise the rhythmic-to-discrete spectrum of arm movements, we considered the arm-pedalling task and the simultaneous electrophysiological recordings described in [9]. In this task, two monkeys (C and D) were trained to control an arm-pedal to move through a virtual environment towards a target (Fig. 1A). Animals performed several types of arm movements that we categorised into 20 conditions: 5 possible numbers of cycles of the arm (*N_cycle_* = 0.5, 1, 2, 4, 7), 2 directions (forward or backwards) and 2 starting positions (top or bottom of the cycle, Fig. 1A-B). Between 1 and 0.5 seconds before movement onset, visual cues indicated to the animal the direction of pedalling and the distance to the target, which was proportional to the number of cycles to be performed. The kinematics of a half-cycle (*N_cycle_* = 0.5; Fig. 1B, blue line) corresponds to a classic discrete movement with distinct start and end positions. In contrast, for seven cycles of pedalling (*N_cycle_* = 7; Fig. 1B, yellow line), the kinematics show a continuous rhythmic movement. Thus, the arm-pedalling task contains an entire rhythmic-to-discrete spectrum of movements. Consistent with this interpretation, simultaneous electrophysiological recordings in SMA, M1, and arm muscles show periodic activity for rhythmic movement (*N_cycle_* = 7, Fig. 1C), a single peak of activity for discrete movement (*N_cycle_* = 0.5, Fig. 1C) and a mix of the two for an intermediate movement (*N_cycle_* = 2, Fig. 1C).

**Figure 1:**
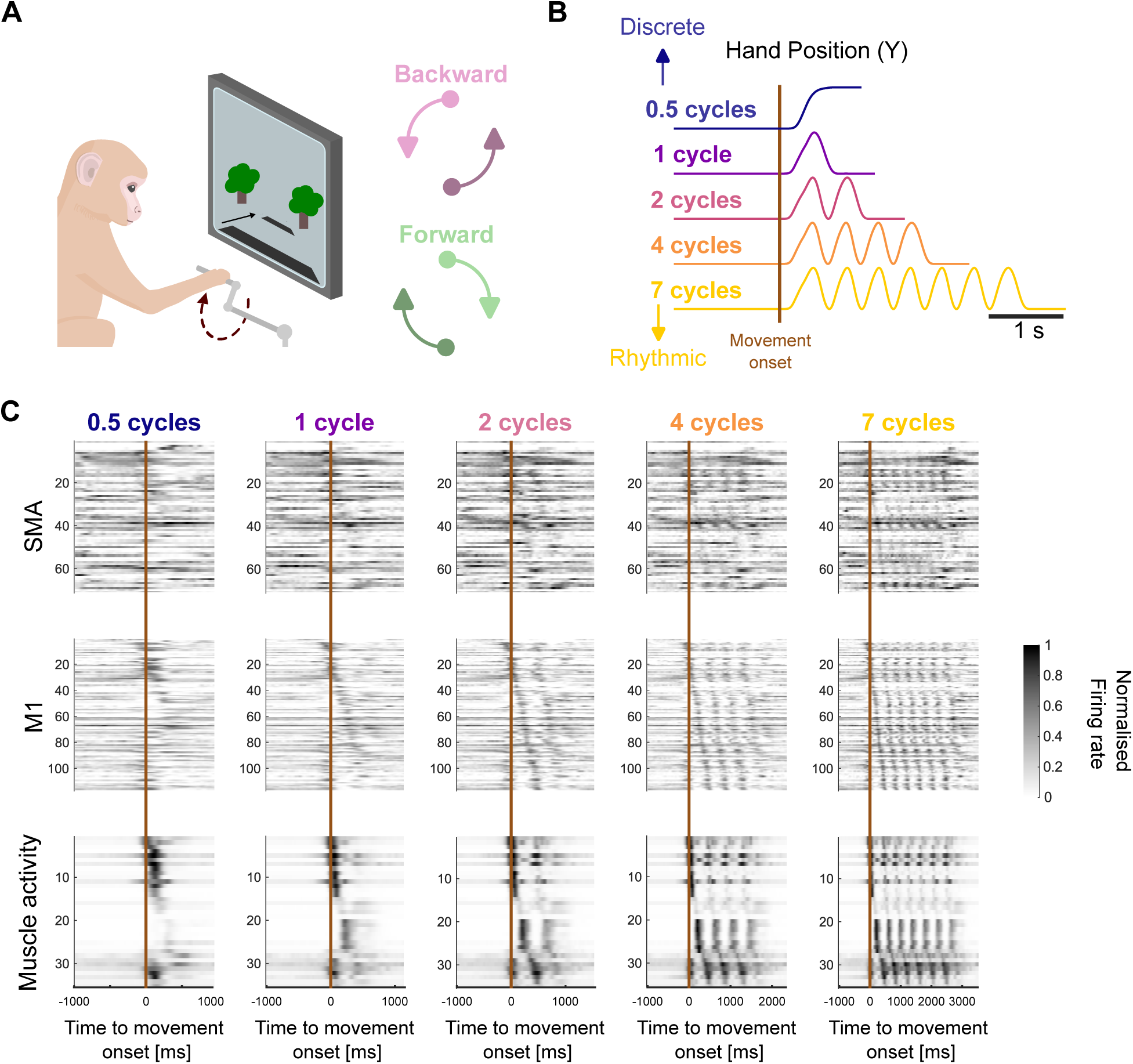
Arm-pedalling task and simultaneous electrophysiological recordings. **A)** Schematic of task. The monkey reached a target on the screen by arm-pedalling through a virtual environment (left). Background colour of the environment instructed the animal to pedal forward/backwards. Distance to the target in the virtual environment determined the number of cycles to be performed (from 0.5 to 7). Pedalling directions and start positions are indicated with different shades in the following figures (right). **B)** Example kinematics from single trials. We plot the hand’s vertical position for each number of arm cycles. **C)** Activity of SMA neurons, M1 neurons, and muscles during forward pedalling, starting from the bottom position. Each row in a heatmap shows the mean firing rate of a unit normalised by its maximum across all conditions. Within heatmaps, rows are sorted by the time of peak activity in the first 500 ms of movement of the 7-cycles condition, so that rows’ identities are comparable across numbers of arm cycles.

### M1 uses the same strategy for rhythmic and discrete movements

We trained RNNs to obtain predictions for M1 dynamics under the hypotheses that it uses the same or different control strategy for rhythmic and discrete movements (Fig. 2A). Both same-control and different-control RNNs were trained to output the low-dimensional trajectories of the muscle activity recorded by EMG while the animals performed the cycling task (Methods). Same-control RNNs received inputs representing the kinematic parameters of the upcoming movement (direction and starting position) and, later, an input to signal to stop the movement (Fig. 2A, left). As a consequence, the only difference between discrete and rhythmic movements was the time when the stop signal arrived. Different-control RNNs received the same inputs as the same-control family, plus a one-hot input indicating the type of upcoming movement (discrete or rhythmic) – in this input encoding, the upcoming movement type was set to 1, the other input to 0. These two inputs entered the network through orthogonal weights.

**Figure 2:**
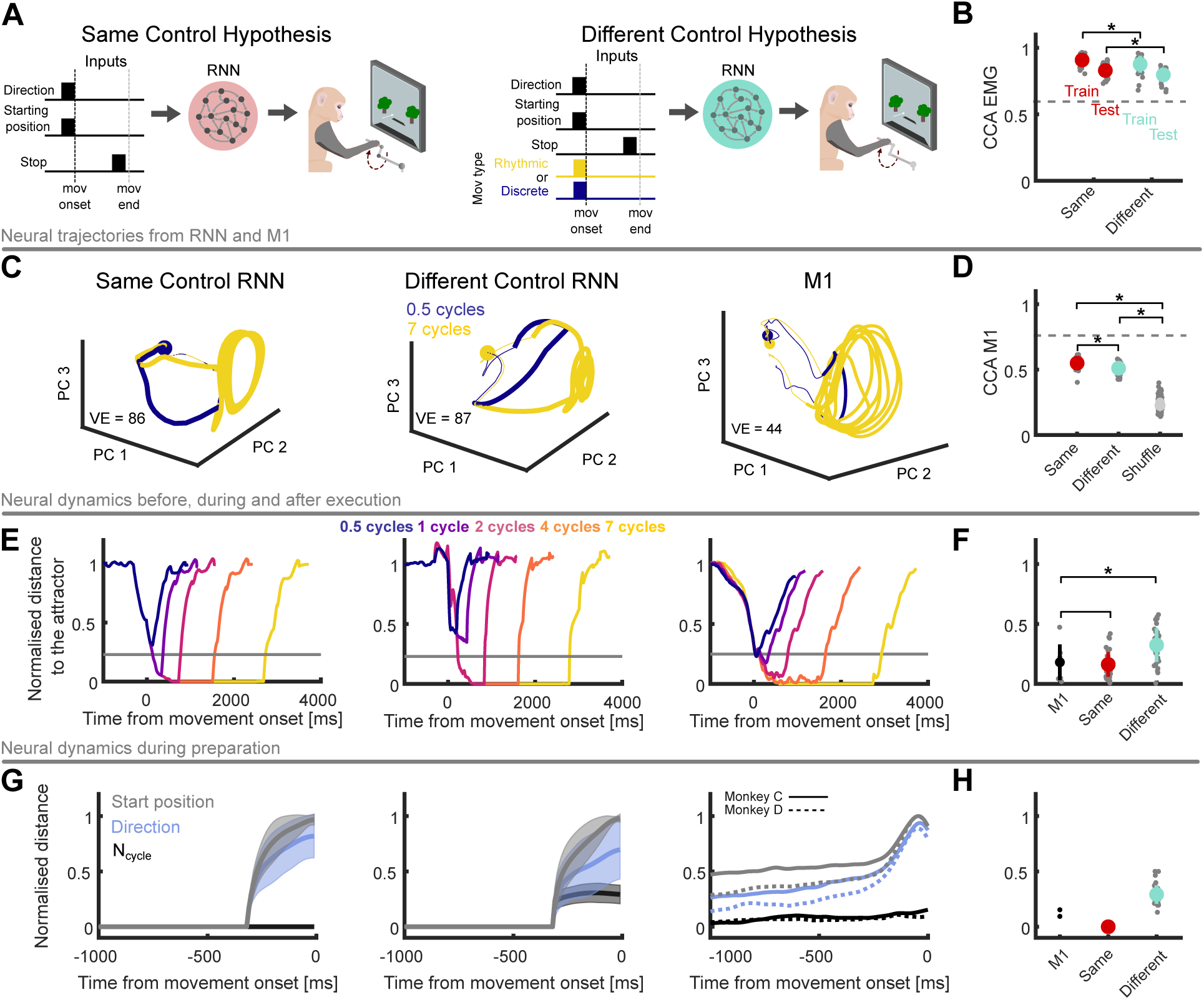
Primary motor cortex shows the same control strategy for rhythmic and discrete movements. **A)** Two families of RNNs were trained to output the low-dimensional trajectories of muscle activity during the task. Left: same control strategy for all movements. Right: Different control strategies for distinct movement types. RNNs received the same inputs as the same-control RNNs, plus an input specifying the type of movement to be performed. **B)** Similarity between EMG trajectories and RNNs’ output measured by Canonical Correlation Analysis (CCA). CC ranges from 0 (uncorrelated) to 1 (identical). Each grey dot corresponds to one RNN. Error bars show standard deviation across RNNs. Dashed line shows CCA across animals as a reference. In all panels, * indicates p-value*<*0.05. **C)** Example neural trajectories of discrete and rhythmic movements. Circles show the start of movement preparation. Thick lines show states during the movement execution. VE: variance explained by the top 3 PCs. **D)** Similarity (CCs) between M1 trajectories and internal dynamics of RNNs. Comparisons were made for M1 versus the trajectories of the trained RNNs and M1 versus the neural trajectories of trained RNNs whose dimensions were shuffled. Error bars show the standard deviation across all RNNs of a family. Dashed line shows CCA across animals as a reference. **E)** Average distance between the neural trajectory and the limit cycle. Distance was 0 when the trajectory was in the limit cycle region and 1 when it was at baseline activity (1s before onset). Average was computed across conditions and animals. All grey lines show the minimum distance for a discrete movement in M1 (blue line). **F)** Minimum distance between neural trajectories of a discrete movement (*N_cycle_* = 0.5) and the limit cycle. Error bars show standard deviation across neural trajectories from different networks and conditions. **G)** Distance during movement preparation between neural trajectories corresponding to different movement features. All distances were normalised by the distance between the neural trajectories at the onset of movements starting in opposite positions. Solid lines, mean across RNNs or conditions (M1); shaded area, standard deviation across RNNs. The M1 panel separately plots data from each monkey. **H)** Distance at movement onset between trajectories corresponding to different numbers of cycles (same direction and starting position). Error bars, standard deviation across RNNs.

We trained 20 RNNs for each family of strategies and each animal (80 in total) to reproduce the EMG trajectories in a subset of the task conditions. Each RNN was trained to output the EMG data in 8 of the 20 conditions. We then tested RNN performance in the 12 held-out conditions. For same-control RNNs, the training set included only rhythmic movement conditions (*N_cycle_* = 4 and 7). For the different-control RNNs, the training set included both ends of the rhythmic-to-discrete spectrum (*N_cycle_* = 7 and 0.5). For both RNN families, the training data comprised the top/bottom and forwards/backwards conditions for the given number of cycles. After training, the RNN outputs from both families successfully reproduced the animals’ EMG activity in both the training and held-out testing sets (Fig. 2B; see Supp. Fig. 1 for example traces). The output of the same-control RNNs showed a marginally but significantly higher similarity with the EMG data than different-control RNNs in both training and testing sets (Fig. 2B, t-test; p-value train = 0.01; p-value test = 5 *·*10^−4^). An exhaustive permutation test confirmed this difference was small: the output of same-control RNNs was more similar to EMG data than the output of different-control RNNs only 65% of the time in training and 68% of the time in testing. Thus, either RNN family could replicate the muscle activity in the training set and correctly inter-or extra-polated to held-out conditions.

The internal dynamics of trained RNNs in both families replicated the main features of M1’s neural dynamics before and during movement (Fig. 2C). Both RNN families predicted that the neural trajectories of rhythmic movements (yellow) would reach a stereotypical rotational pattern – a limit cycle – during movement execution (we quantify the limit cycle in Supp. Fig. 2A). Both families also predicted that neural trajectories of discrete movement (blue) approached and quickly receded from the limit cycle, thus also showing rotational dynamics (Supp. Video 1). We found the same dynamics in M1 (Fig. 2C). Notably, in M1 and both RNN families, all neural trajectories approached the limit cycle on a plane orthogonal to the limit cycle itself; thus, neural trajectories of discrete movements and rhythmic movements rotated on orthogonal planes (Supp. Fig. 2B-C).

Both RNN families’ predictions for the neural trajectories in held-out conditions were more similar to those of M1 than were shuffled controls (t-test, p-value *<* 10^−10^ for both families). Permutation tests confirmed that both RNN families’ trajectories were more similar to M1 than shuffled controls 100% of the time. The internal dynamics of same-control RNNs showed a slightly, but significantly higher similarity to M1’s dynamics than different-control RNNs (Fig. 2D, t-test p-value = 2 *·* 10^−4^). Exhaustive permutation tests showed same-control RNNs were more similar to M1 trajectories than were different-control RNNs 75% of the time. These results confirmed that either the same-or different-control hypothesis is a plausible solution to how M1 controls the rhythmic-to-discrete spectrum of arm movements.

To further test the validity of both models of control, we trained a set of 20 “unconstrained” networks for each animal (40 total) in which the weights of the inputs encoding the rhythmic and discrete movements were also optimised during training. If both the same- and different-control hypotheses are plausible models of cortical dynamics, as our results above suggest, then training networks without any assumption about the separation of the input weights should yield networks that converge on either solution. That is what we found (Supp. Fig. 3): across the 40 networks, we found some for which the rhythmic and discrete movement inputs were weighted identically, like our same-control RNNs, some for which those inputs had orthogonal weights, like our different-control RNNs, and others with intermediate angles between the input weights (average angle between input weight vectors was 37*^◦^*). This result shows that unconstrained optimisation of the input weights is not sufficient to determine whether the same-or different-control hypothesis is in place in M1. Instead, it is crucial to compare the internal dynamics of the trained networks to the dynamics of M1 to disentangle the hypotheses.

The preceding results also showed that solely observing cortical population dynamics cannot differentiate the hypotheses of same or different control strategies. Previous work has established that M1 shows latent rotational dynamics during both discrete and rhythmic movements [8–10, 14]. Yet, as we show here, both families of RNNs showed rotational dynamics during the execution of discrete and rhythmic movements, so observing them in M1 during both movement types cannot differentiate the same- and different-control hypotheses. Similarly, while the separation of neural trajectories into different subspaces has been proposed as evidence of different control strategies [12, 13], both our families of RNNs show this separation of neural trajectories into orthogonal subspaces (Supp. Fig. 2B-C), so observing this separation in M1 during both movement types also cannot differentiate the two hypotheses.

Our RNN models predicted that what can differentiate the hypotheses is how closely neural trajectories approach the limit cycle during discrete movements (See Fig. 2C and Supp. Fig.2B for examples). Same-control RNNs predicted that neural trajectories smoothly approach the limit cycle during discrete movements, coming closer to the limit cycle the closer the movement is to being fully rhythmic (Fig. 2E, left). Different-control RNNs predicted that the neural trajectories of discrete movements always extend the same, shorter distance towards the limit cycle (Fig. 2E, middle). Neural trajectories in M1 matched those of the same-control RNNs (Fig. 2E, right), smoothly approaching the limit cycle during discrete movements (Fig. 2F, t-test (M1, same-control), p-value = 0.67, t-test (M1, different-control), p-value = 0.01).

The two families of RNNs also made distinct predictions for the dynamics of M1 during movement preparation, in the second leading up to movement onset (Fig. 2G; see Supp. Fig. 4 for examples). Same-control RNNs predicted that, during preparation, neural trajectories would be the same for both upcoming discrete and rhythmic movements, but would differ if the direction or starting position of movement varied (Fig. 2G, left). By contrast, different-control RNNs predicted that, during preparation, neural trajectories would separate according to the type of upcoming movement (Fig. 2G, middle). Neural trajectories of M1 matched those of same-control RNNs, showing a minimal separation for trajectories of upcoming movements of different numbers of cycles (Fig. 2G; Supp. Fig. 4). At the time of movement onset, M1’s trajectories corresponding to different numbers of cycle were closer to the normalised distance predicted by same-control RNNs than by different-control RNNs (Fig. 2H; absolute difference of averages: (M1-Same) = 0.12, (M1-Different) = 0.17).

To rule out the possibility that the neural trajectories of M1 separated by movement type in any dimension, we used demixed Principal Component Analysis (dPCA, [15]) to identify axes that encoded the number-of-cycles in both families of RNNs and in M1 activity. Same-control RNNs predicted that any identified number-of-cycles axis would not separate neural trajectories and would explain less variance of activity than the axes encoding the start position, direction of cycling, and condition-independent dynamics (Supp. Fig. 5). Conversely, different-control RNNs predicted the number-of-cycles axis would separate neural trajectories and explain more variance of activity than the axis encoding the direction of movement. The identified number-of-cycles axes in M1 activity matched the predictions of the same-control RNNs. Thus, across our analyses, M1 neural trajectories consistently matched the predicted dynamics of networks that employ the same control strategy across the spectrum of discrete and rhythmic movements.

We considered the possibility that the poorer predictions of M1 dynamics by different-control models arose from how we chose to model their contextual inputs. To test this, we trained two variants of the different-control RNNs (Supp. Fig. 6). As for the RNNs in Fig. 2A, both variants received inputs indicating the direction and starting position of the upcoming movement. The key difference was in the encoding of the type of movement. The first variant received the same inputs as the different-control model in Fig. 2A for rhythmic movements. But for discrete movements, it did not receive a stop signal: instead, the number of cycles (0.5 or 1) was encoded in the height of the input (Supp. Fig. 6A). A second variant encoded the discrete movements in the same way as the first, but encoded the rhythmic movements with a constant input that lasted until 100 ms before the movement’s end (Supp. Fig. 6B). Networks of the first variant struggled to solve the task (Supp. Fig. 6C, left). Networks of both families produced internal dynamics less similar to M1 dynamics than the original models (Supp. Fig. 6C, right). This effect was observed even when the variants were trained for longer than the original networks (30000 epochs for the originals; 50000 epochs for these variants). Networks of both families that were considered successful did not recover the main internal dynamics of M1, either showing rotational trajectories for discrete movements, or rotations decaying with every cycle of rhythmic movement (Supp. Fig. 6A-B, right). Consequently, our results were not specific to the kind of contextual input we used for the different-control networks.

### SMA uses different strategies for rhythmic and discrete movements

SMA is a key region for the production of rhythmic movements [9, 11] and sequences of discrete movements [16]. Hence, we next asked whether SMA used the same or a different control strategy for both movement types. We again trained two families of RNNs, each embodying a control hypothesis (Fig. 3A). Because SMA densely projects to M1 [17], we trained the RNNs to output the low-dimensional neural trajectories of M1. Training and test conditions were the same as in the M1 models. Just as in our M1 models, the RNNs modelling SMA received the direction and starting position of the upcoming arm movement. Unlike RNNs modelling M1, RNNs modelling SMA received a temporal-context input throughout the movement, which allowed them to track duration of movement, and was necessary to replicate the characteristic helical dynamics of SMA during this task [9] (Supp. Fig. 7). The temporal-context input was constant from 800 to 300 ms before movement onset and then linearly decreased up to the movement offset (Fig. 3A, left). In the case of the different-control RNNs, the temporal-context signal was a one-hot input for discrete and rhythmic movements that entered the networks through orthogonal sets of weights (Fig. 3A, right).

**Figure 3:**
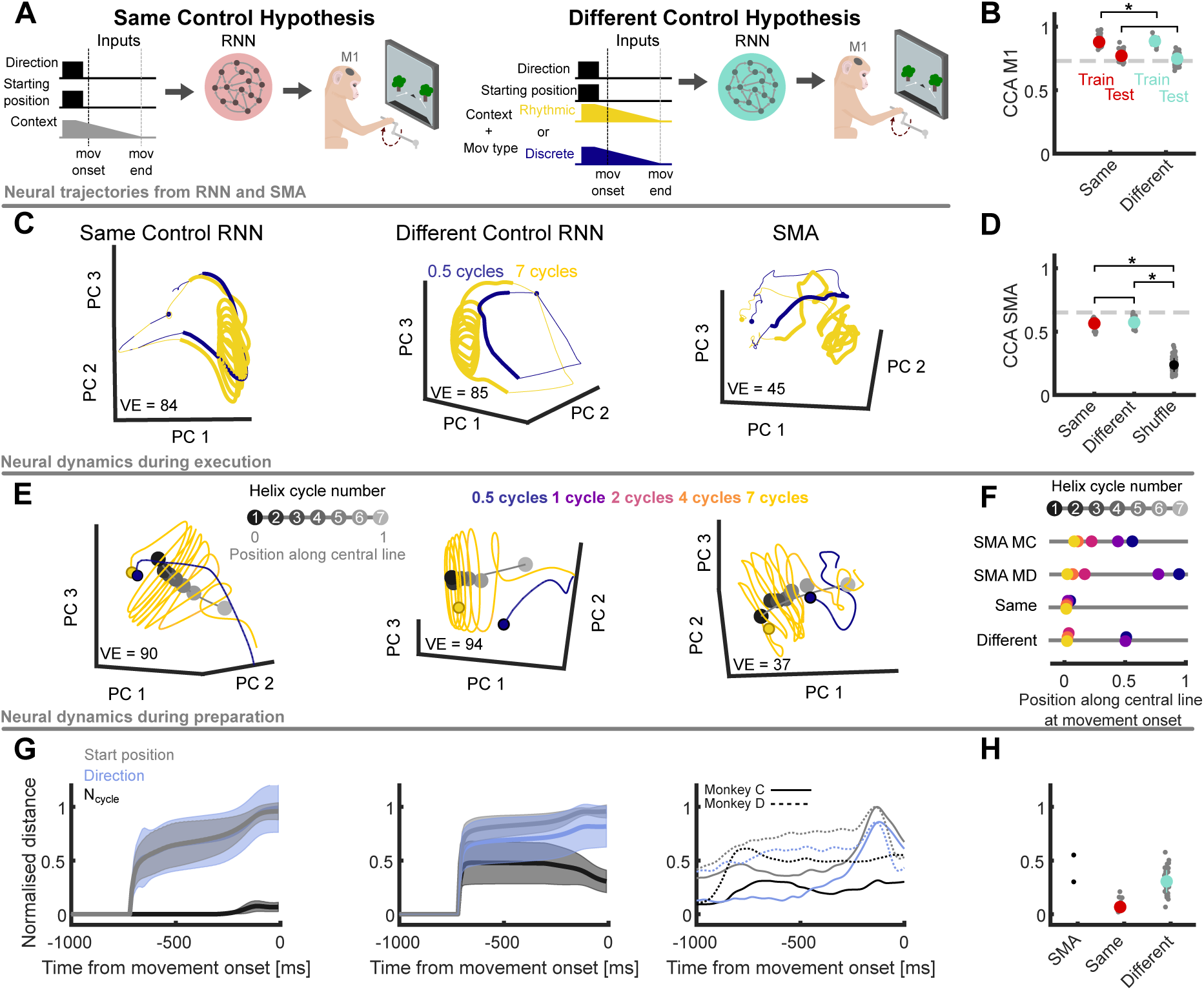
Supplementary Motor Area shows different control solutions for rhythmic and discrete movements. **A)** Two families of RNNs were trained to output the low-dimensional activity of M1 during the task. Left: same control strategies for all movements. Right: different control strategies for distinct movements. RNNs received the same inputs as the same-control RNNs, but the temporal-context input specified the type of movement (discrete or rhythmic) **B)** Similarity (CCs) between M1 trajectories and RNNs’ output. Each grey dot corresponds to one RNN. Error bars show the standard deviation across RNNs. Dashed line shows CCA across animals as a reference. In all panels, * indicates p-value*<*0.05 **C)** Example neural trajectories from discrete and rhythmic movements. All trajectories correspond to pedalling forward from the bottom. Circles show the start of movement preparation. Thick lines show states during the movement execution. VE shows the variance explained by the top 3 PCs. **D)** Similarity (CCs) between SMA trajectories and internal dynamics of RNNs. Comparisons were made for SMA versus the dynamics of the trained RNNs and SMA versus the dynamics of trained RNNs whose projections into neural dimensions were shuffled. Dashed line shows CCA across animals as a reference. **E)** Example neural trajectories during the execution of rhythmic and discrete movements. Coloured circles indicate the trajectories’ positions at movement onset for different numbers of cycles. Black to grey circles show the average activity for each helix cycle of the 7-cycle movement. Grey line shows the central line of the helix, which was fitted to the centres of the cycles. **F)** Average position of the neural trajectories of different number of cycles at movement onset along the central line of the helix. Average was computed across conditions for each animal. Vertical position of each dot was randomised around the grey line for visualisation purposes only. **G)** Distance during preparation between neural trajectories corresponding to different movement features. All distances were normalised by the distance between the neural trajectories at the onset of movements that started in opposite positions (same direction and number of cycles). Solid lines, mean across RNNs or conditions (SMA); shaded area, standard deviation across RNNs. The SMA panel separately plots data from each monkey. **H)** Normalised distance between trajectories corresponding to different number of cycles at movement onset.

Both families of trained RNNs successfully output M1’s population dynamics during pedalling movements (Fig. 3B). The output of different-control RNNs showed a marginally but significantly stronger similarity to M1’s dynamics than did same-control RNNs on the training set but not the testing set (Fig. 3B, t-test: p-value train = 0.027, p-value test = 0.31; permutation test was consistent: different-control outperformed same-control in 63% of cases in training, 38% in testing). Both families’ internal dynamics showed the key features of SMA’s dynamics (Fig. 3C) [9], traversing a helical trajectory around a central axis from the first to the last cycle of a rhythmic movement. In both families of RNNs and in the SMA, the trajectory rotated around the helix’s central axis as many times as there were cycles of the arm being produced (Supp. Video 2). Again, in held-out conditions, the RNNs’ predicted neural trajectories well matched those from the SMA (Fig. 3D). The internal dynamics of same- and different-control RNNs showed no significant difference in their similarity to SMA’s dynamics (Fig. 3D, t-test p-value = 0.29; permutation test: different-control outperformed same-control RNNs in 54% of cases). Both RNN families’ predictions for the neural trajectories in held-out conditions were more similar to those of SMA than were shuffled controls (t-test, p-value *<* 10^−10^ for both families; permutation test: both families of RNNs outperformed shuffled RNNs in 100% of cases). Similar to the case of M1, training “unconstrained” networks without fixed input weights yielded a set of networks that showed the entire spectrum of possible input-weight solutions (Supp. Fig. 3B,C, average angle between inputs = 44*^◦^*).

Both RNN families predicted that, at the end of movement, the trajectories converged to the same end position along the helix’s central axis, regardless of the number of cycles performed. SMA’s trajectories also converged at the end of movement (Fig. 3E). The two RNN families made different predictions for the initial conditions of the SMA’s trajectory at the onset of discrete and rhythmic movements. Same-control RNNs predicted that, at movement onset, neural trajectories start at the same position along the helix’s central axis regardless of the number of cycles to be performed (coloured circles in Fig. 3E, left). Consequently, during execution, their trajectories cover the same distance along the helix’s central axis, their paths differing by the number of cycles performed. Different-control RNNs predicted that, at movement onset, neural trajectories start at different positions along the central axis of the helix (Fig. 3E, middle). Notably, they predict the starting positions cluster by the type of movement (Fig. 3F, bottom). Consequently, during execution, the trajectories cover different distances along the helix’s central axis, and their paths differ by the number of cycles performed according to whether the movement is rhythmic or discrete. SMA’s neural trajectories more closely matched those of different-control RNNs (Fig. 3E, right), with the starting positions for discrete movements clearly separated along the central axis of the helix from the starting positions for rhythmic movements (Fig. 3F).

That SMA’s neural trajectories have different initial conditions at movement onset implies that they must diverge during the preceding period of movement preparation. We thus checked the RNN families’ predictions for movement preparation. The same-control RNNs predicted that neural trajectories during preparation would not separate according to the number of cycles (Fig. 3G, left), but different-control RNNs predicted that they would (Fig. 3G, middle), and would separate as much as trajectories corresponding to different directions and starting positions (Supp. Fig. 8). SMA trajectories during movement preparation again matched the predictions of different-control RNNs (Fig. 3G). At the time of movement onset, the average normalised distance between SMA’s trajectories corresponding to different number of cycles was closer to the normalised distance shown by different-control RNNs than by same-control RNNs (Fig. 3H; absolute difference of averages: (SMA-Same) = 0.36, (SMA-Different) = 0.12). To assess the extent of the influence of the number of upcoming movement cycles on the dynamics of SMA, we again used dPCA to find encoding axes in RNN and SMA activity (Supp. Fig. 9).

The number-of-cycles axes found by dPCA explained as much variance as direction and starting position axes in different-control RNNs and SMA, but explained less variance in the same-control RNNs (Supp. Fig. 9B), again consistent with SMA using different control strategies for discrete and rhythmic movements.

In addition to M1, SMA also projects to the spinal cord [18]. To test if our results depended on the SMA’s downstream target, we trained the same and different-control networks for SMA to directly output EMG trajectories. Networks successfully learned the task and similarly showed that SMA’s dynamics matched the predictions of different-control RNNs (Supp. Fig. 10). Thus, our inference that SMA uses different strategies for controlling discrete and rhythmic movements is robust to the choice of downstream target for SMA activity.

We did though observe a disagreement between the predictions of the different-control RNNs and the activity of SMA during preparation. The different-control RNNs predicted only two possible initial conditions at movement onset: one for rhythmic movements, and one for discrete movements (Fig. 3F; Supp. Fig. 9). By contrast, SMA’s population activity had a range of initial conditions at movement onset within the groups of rhythmic and discrete movements (Fig. 3F, top). Moreover, those conditions were ordered by the number of upcoming cycles, consistent with our dPCA observation that neural trajectories during preparation were sorted along a number-of-cycles axis (Supp. Fig. 9).

This coding of the number of cycles implies SMA receives input that not only differentiates rhythmic and discrete movements, but also encodes the number of cycles of those movements. We thus tested already-trained different-control RNNs with a different input design, one in which the extremes of discrete (0.5 cycle) and rhythmic (7 cycle) movements were again encoded by orthogonal input weights, but intermediate movements (1, 2, 4 cycles) were encoded by rotating the weight vector between these two extremes (Fig. 4A). This rotating-input RNN still reproduced SMA’s helical trajectories during movement execution (Fig. 4B). Unlike the original different-control RNNs, it did so while also creating initial conditions at movement onset that were grouped by movement type, yet ordered by the number of upcoming cycles (Fig. 4C, bottom). Consequently, as in SMA, neural trajectories of the rotating-input RNNs separated according to the upcoming number of cycles (Fig. 4D) and were sorted along a number-of-cycles encoding axis (Supp. Fig. 11E). Further, also like SMA, the rotating-input RNNs’ activity had axes encoding cycle number that explained as much variance as the axes encoding direction (Supp. Fig. 11F). Consequently, the rotating-input RNN suggests SMA has elements of both different- and same-control strategies for arm movement: distinct representations of rhythmic and strongly discrete arm movements, but a continuum within each type of movement.

**Figure 4:**
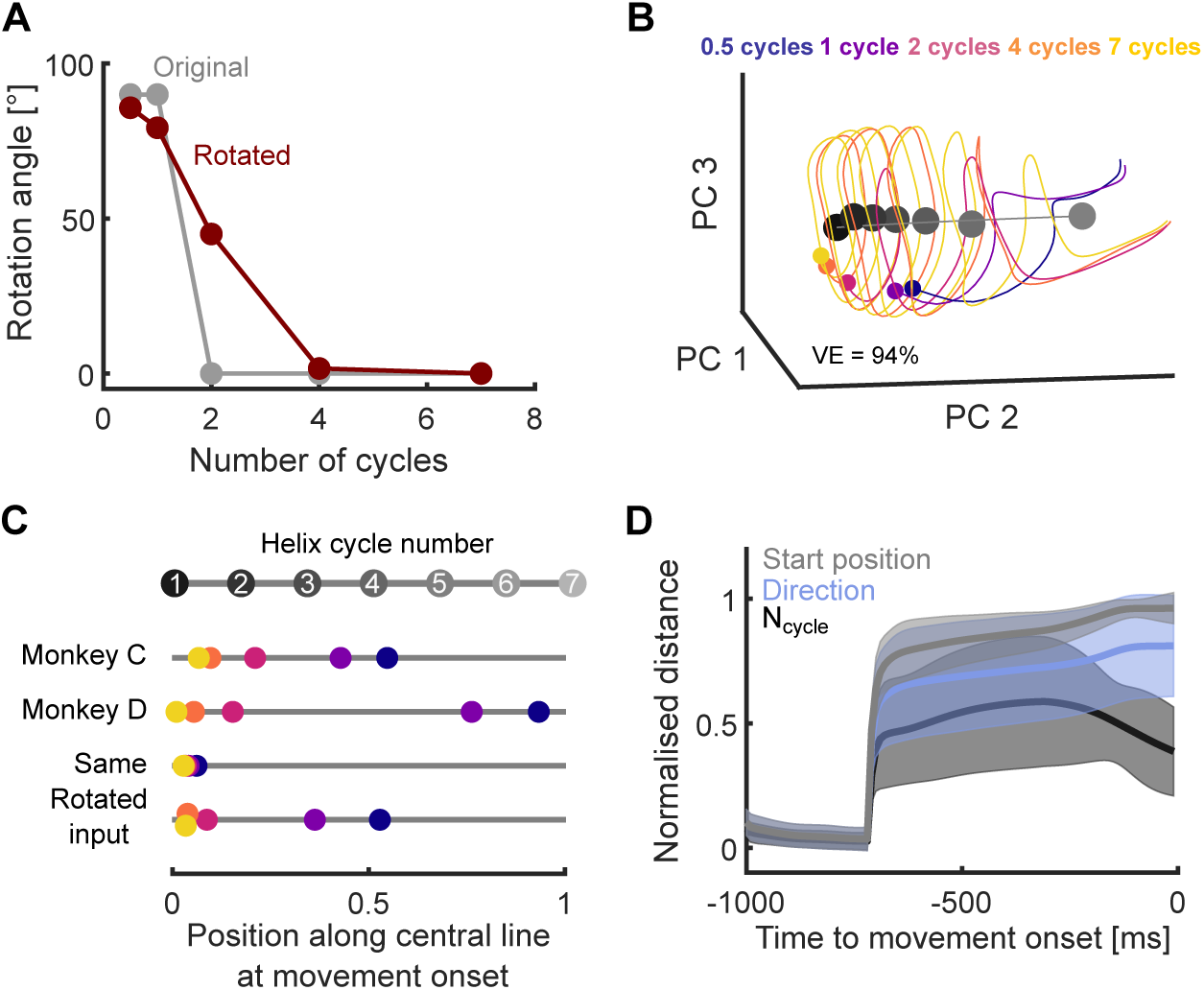
Different-control RNNs with rotating input match SMA neural dynamics. **A)** Angle of the temporal-context input with respect to the 7-cycles condition. Original (different-control) RNNs assumed that inputs of rhythmic and discrete movements were orthogonal. **B)** Rotating-input RNNs produce helical trajectories during movement execution. VE: variance explained by top 3 PCs. **C)** Average starting position of neural trajectories at movement onset for rotating-input RNNs, same-control RNNs, and SMA activity. **D)** Distance during preparation between rotating-input RNN neural trajectories corresponding to different movement features. Compare to Fig. 3G, right.

## 3 Discussion

The possible cortical solutions that give rise to the control of rhythmic and discrete movements have been debated for decades [1]. Two seemingly opposing hypotheses have been proposed: these types of movements emerge from the same or different cortical control strategies. To test which hypothesis could account for cortex’s solution to controlling movements across the rhythmic-to-discrete spectrum, we exploited the comparable kinematics across rhythmic and discrete movements in an arm-pedalling task and the modelling power of RNNs. The arm-pedalling task allowed us to examine neural dynamics during the complete rhythmic-to-discrete spectrum of movements, while the RNNs allowed us to make predictions of the neural dynamics under each hypothesis of control. Our results reconcile these apparently opposite views by showing evidence that both solutions are implemented in different motor cortical areas. While M1 employs the same strategy for controlling both movement types, SMA, an area that contributes to higher-order aspects of motor control, exhibits different strategies for the two types of movements.

Our results propose parsimonious explanations for puzzling observations regarding the dynamics of M1 and SMA. M1 shows strongly rotational dynamics during rhythmic [9, 10, 19] and discrete movements [8, 14, 20]. While the rotational dynamics could drive the kinematics of rhythmic movements, its computational role during discrete movements is as yet unclear [8, 20–23]. By directly contrasting M1 population dynamics for discrete and rhythmic movements, our results offer a simple explanation for the rotational dynamics of M1 during discrete movements: they are a consequence of the neural trajectories approaching and then smoothly returning from the limit cycle (Fig. 2). Thus, M1’s strategy for generating a movement – regardless of its rhythmicity – is to drive neural activity to the limit cycle and abandon it to end the movement.

In the case of SMA, recent evidence shows that it has strongly helical dynamics during rhythmic movements [9], and predominantly non-rotational dynamics during discrete movements [11]. As these rhythmic and discrete movements were studied in separate tasks, it was unclear if these dynamics emerged from the same strategy for movement control. We find SMA has a single dynamical solution for all movements: all neural trajectories follow a helix that unfolds for as many rotations as the number of cycles performed (Fig. 3C-E). Yet, the strategies to control rhythmic and discrete movements are different. Neural trajectories of rhythmic and discrete movements diverge during their preparation to reach different positions along the central axis of the helix at movement onset. The more rhythmic the movement, the further away from the end point of the helix. Consequently, the neural trajectories of discrete movements begin near the end point of the helix at movement onset and only perform a brief turn during execution, thus explaining why previous reports have seen less rotational dynamics in SMA during discrete than rhythmic movements [11].

While SMA dynamics were better-matched by different-control models, they also showed features consistent with both models of control. Specifically, at movement onset, neural trajectories of SMA were separated by movement type, yet within those types, trajectories were sorted by the upcoming number of cycles (Fig. 3F). In addition, before movement onset, neural trajectories separated by the upcoming number of cycles (Supp. Fig. 9), suggesting some continuity within the rhythmic-to-discrete spectrum of movements. We could capture these features by extending the different-control model to include a continuous encoding of the number of cycles for movements intermediate between strongly discrete and strongly rhythmic. Thus, our results suggest the SMA’s control of discrete and rhythmic arm movements is more ambiguous than the dichotomy of same-or different-control.

Our results propose a new model for the roles that M1 and SMA play in controlling the rhythmic-to-discrete spectrum of arm movements. The model comprises the following sequence of events. First, during movement preparation, neural trajectories of SMA diverge according to the upcoming number of movement cycles. This divergence is not transmitted to M1, whose preparatory dynamics do not encode the movement type. Second, at movement onset, SMA’s trajectories fall onto a helical spiral. Third, during movement execution, SMA’s neural activity traversing the helix from that starting point to the end of the helix encodes the number of cycles of arm movement. Traversing only a single cycle or less of the helix creates a discrete movement. At the same time, in M1, neural trajectories move toward a limit cycle: for discrete movements, a stop signal prevents the trajectory from reaching the limit cycle; for rhythmic movements, the trajectory reaches and traverses the limit cycle until a stop signal arrives. That stop signal could originate from SMA: that all SMA trajectories for discrete and rhythmic movements converge on the same end position is consistent with that end position signalling the end of movement.

This model is congruent with the evolutionary history of the motor cortex. It has long been proposed that cortex’s fine control of discrete limb movements is built on its existing dynamics for controlling evolutionarily older rhythmic movements such as locomotion [24]. M1 appeared earlier in the evolutionary history of mammals than the SMA [25, 26]. Our finding of a limit cycle in M1 is consistent with it being the evolutionarily older structure driving rhythmic behaviour, including locomotion, in early mammals. Further, our data and model suggest SMA’s dynamics provide a timing signal for the production of discrete movements and the duration of rhythmic movements. This is consistent with the later evolution of SMA providing top-down signals to M1’s existing dynamics that allow fine limb control, and is consistent with the increased behavioural repertoire associated with its evolution [26]. The evolutionary timeline of the fine control of limb movement is thus one possible answer to the question of why SMA and M1 have different cortical strategies for the spectrum of rhythmic-to-discrete movements.

Our model provides two hypotheses for the control of arm movements that are yet to be explored experimentally. First, the model proposes that the control of discrete and rhythmic movements by M1 stems from a limit-cycle attractor. It follows that driving the neural activity of M1 to or away from the limit cycle should respectively start and end movements with highly precise timing. Second, our model also proposes that SMA drives the timing of the inputs to M1 that end muscle activity. Consequently, perturbing the dynamics of SMA should hinder the control of the timing of discrete and rhythmic movements. Pushing SMA activity from the helical spiral should move M1 activity off its limit cycle, interfering with muscle activity. Probing these predictions experimentally would provide further, causal evidence for the parallel control of rhythmic and discrete movement in motor cortex.

## 4 Methods

### 4.1 Behavioural task

Here we considered an arm-pedalling task described in detail in [19]. Briefly, two rhesus macaque monkeys(*Macaca mulatta*, C and D) were trained to go through a virtual environment by moving a pedal with their right arm. To minimise movement of the opposite arm, the left arm was loosely restrained during the task. Animals were instructed to pedal forward or backwards as indicated by the landscape colour of the virtual environment. Monkeys pedalled on the sagittal plane and started from either the top or the bottom of the wheel. In each trial, the animals performed a varying number of cycles (0.5, 1, 2, 4 or 7) to reach an objective. Visual cues indicating the movement direction and the number of cycles to be performed were provided to the animal 0.5 to 1 s before the movement started. The background colour of the environment indicated the pedalling direction (green = forward, orange = backwards). The distance to the objective in the virtual environment was proportional to the number of cycles. After the animal reached the objective, its arm must remain stationary to receive a juice reward. This way, each trial comprised three epochs: preparation with the arm stationary (for 1 to 1.5 s), cycling (each cycle lasted *≈* 500 ms) and a stop with the arm stationary (1.5 s). Each movement was classified by its direction, starting position and number of cycles, resulting in 20 different conditions (2 directions *×* 2 start positions *×* 5 number of cycles). Monkeys performed all combinations of two cycling directions, two starting orientations, and five distances. Trials were grouped into forward- and backward-cycling blocks. Other trial types were randomly interleaved within those blocks.

A trial was considered successful if it met the following conditions: the animal started cycling within 650 ms after the go cue; once cycling began, the final target had to be reached within a distance-dependent time limit; the speed was above a threshold at all times during the cycling. If the monkey moved past the final target or moved after acquiring the target, the trial was aborted. These constraints encourage stereotypical movements across trials. Approximately 15 trials per condition were collected for both monkeys.

### 4.2 Neural recordings

A detailed description of neural recordings can be found in [9]. Briefly, neural recordings were obtained using conventional single electrodes. Single neurons were recorded consecutively from SMA and M1. Motor cortex recordings were restricted to regions where microstimulation elicited responses in the shoulder, upper arm, chest, and forearm. Nearly all neurons that could be isolated in motor cortex were responsive during cycling. A small number of neurons were discarded due to low signal-to-noise ratios or insufficient trial counts. No further selection criteria were applied. SMA and M1 recordings contained between 71-77 and 116-117 neurons, respectively.

Intra-muscular EMG was recorded from the major muscles of the arm, shoulder, and chest using percutaneous pairs of hook-wire electrodes. A total of 27-35 sites were recorded. Recordings were made from 1-8 muscles at a time, on separate days from neural recordings. Recordings were made from the following muscles: the three heads of the *deltoid*, the two heads of the *biceps brachii*, the three heads of the *triceps brachii, trapezius, latissimus dorsi, pectoralis, brachioradialis, extensor carpi ulnaris, extensor carpi radialis, flexor carpi ulnaris, flexor carpi radialis,* and *pronator*.

### 4.3 Defining neural trajectories

Principal Component Analysis (PCA) was applied to extract the low-dimensional activity of the neural and muscle population. Recordings were preprocessed as follows. Neural activity was selected from 1 second before the movement onset to 400 ms after the movement ended. Spike trains were smoothed using a Gaussian filter of standard deviation *σ* = 25 ms. The average firing rate of each neuron was soft-normalised by dividing by a normalisation factor (normalisation factor = firing rate range + 5) [9, 19]. PCA was performed across all conditions’ averaged neural activity. Neural trajectories were defined as the projection of the firing rates of the populations onto the Principal Components (PCs). Trajectories had as many dimensions (*N_d_*) as necessary to explain at least 80% of the variance unless stated otherwise. Neural trajectories were recentred so that the origin of the subspace corresponded to the baseline activity of the population before the movement preparation started. We defined the baseline activity as the average neural activity across conditions at 1 s before movement onset. This way, the origin of the subspace across recordings is a common physiological reference (Fig. 2, Fig. 3, Supp. Fig. 2). The same process was followed to define low-dimensional muscle trajectories.

### 4.4 Recurrent Neural Networks (RNNs)

#### 4.4.1 M1

We trained two families of RNNs to solve the arm-pedalling task. Both types of networks consisted of N = 50 firing-rate units, and their target outputs were the low-dimensional trajectories (*N_d_* = 4) of the muscle activity (EMG) during the task. These dimensions explained 78 and 88% of the variance of EMG recordings from monkey C and D, respectively. The first family (same-control RNNs) assumed that all movements were generated from the same control strategy. Then, the only difference between the number of cycles performed stemmed from variations in the timing of the inputs that trigger the end of arm movements. The dynamics of a network of this family was given by:

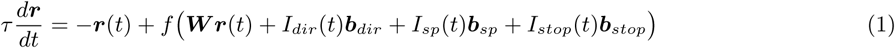

where *τ* is a time-constant, ***r*** represents an N-dimensional vector of firing rates, *f* = *tanh* is a nonlinear input-output function, ***W*** is an N by N matrix of recurrent weights. Considering that the neural and EMG recordings were smoothed using a Gaussian filter (*σ* = 25 ms) and subsequently trial averaged, *τ* was set to 40 ms for all networks. *I_dir_*, *I_sp_* and *I_stop_* represent time-varying external input indicating the cycling direction, starting position and moment to stop, respectively. All inputs were square with an amplitude equal to one and lasted 300 ms. *I_dir_*, *I_sp_* ended at the time of movement onset, while *I_stop_* ended 100 ms before the end of the movement. ***b****_dir_*, ***b****_sp_*and ***b****_stop_*are vectors corresponding to the projection weights of each input into the network. ***b****_stop_* is a vector of parameters shared across all conditions. The input pulse corresponding to a movement in a specific direction (*I_dir_*) entered the network through different sets of random input weights depending on the direction of he upcoming movement: ***b****_dir,backward_* and ***b****_dir,forward_*. The same strategy was applied to model the start position. The output of the network was

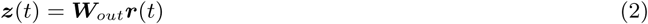

where ***z*** is a 4-element vector corresponding to the output of the network, while ***W****_out_* is a 4*×*N weight matrix. Networks were trained using back-propagation-through-time using TensorFlow and an Adam optimiser to adjust ***W***, ***b****_dir_*, ***b****_sp_*, ***b****_stop_*, ***W****_out_* to minimise the squared difference between the network output (***z***) and the low-dimensional muscle activity. The initial values of the input weights were drawn from a zero-mean normal distribution (*σ* = 0.3) and adjusted throughout training. We considered a time step Δ*t* = 10 ms for all networks. Crucially, this family of networks was trained only using the conditions of rhythmic movements (*N_cycle_* = 4 and *N_cycle_*= 7, all directions and starting positions), so that the dynamics of the RNNs for discrete movements (*N_cycle_* = 0.5 and *N_cycle_*= 1) were a *de novo* prediction of the dynamics of M1 for these conditions. All networks were trained for 30,000 epochs. Networks were considered successfully trained if the Pearson correlation between their output and the EMG low-dimensional trajectories was at least 0.8 across train and test conditions. Networks that were not considered successful (3 for same-control, 1 for different-control) were discarded from subsequent analyses.

The second family of networks (different-control RNNs) assumed that discrete and rhythmic movements were generated by different control strategies. We modelled this assumption as an additional input indicating the kind of movement to be generated (*I_mov_*). Therefore, the dynamics of these networks was:

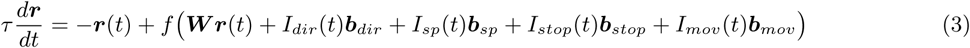

Where ***b****_mov_* is a vector of weights that project the input *I_mov_* into the network. ***b****_mov_* could take two forms depending on the kind of movement to be generated: ***b****_mov,discrete_* or ***b****_mov,rhythmic_*. The timing and magnitude of *I_mov_* was identical to the input of direction and starting position. To encourage the network to find different solutions for different kinds of movements, we defined ***b****_mov,discrete_* and ***b****_mov,rhythmic_* as two orthogonal vectors drawn from a normal distribution before training. These vectors were not modified during training. This family of networks were trained with all directions and starting positions of the conditions *N_cycle_*= 0.5 (discrete) and *N_cycle_* = 7 (rhythmic). RNN dynamics were then predicted for all remaining conditions. Networks trained to output EMG data of monkey C and D were trained for 50,000 and 30,000 epochs, respectively.

We also trained “unconstrained” RNNs in which the sets of weights of the movement type inputs were also optimised by the training of the networks (Supp. Fig. 3). These networks used the same inputs to specify direction, starting position and stop signal as defined above. The weights (***b****_mov,discrete_* and ***b****_mov,rhythmic_*) were initialised by sampling from a uniform distribution over the interval [*−*0.05, 0.05]. RNNs were trained for 50,000 epochs.

Variants of the two models (Supp. Fig. 6) used the same inputs described above to specify direction and start position. Two variants of the different-control model were proposed. For the first variant, the networks did not receive a stop signal for discrete movements; instead, the number of cycles to be performed (half or 1 cycle) was encoded in the height of the discrete input (Supp. Fig. 6A). A second variant encoded the rhythmic movements with a constant input from 300 ms before movement onset until 100 ms before the end of the movement (Supp. Fig. 6B). For both of these variants, the sets of weights corresponding to the movement-type inputs were orthogonal. Unlike the original models, variants of different-control models were trained using conditions of *N_cycles_* = 0.5, 1, 4 and 7 cycles so that the networks were trained with examples of two numbers of cycles for each movement type. Preliminary tests showed that most variants trained with the same number of epochs as the original models were not successful. For this reason, all variants were trained for 50,000 epochs, two-thirds more than the original same-control model.

#### 4.4.2 SMA

We trained two families of RNNs to output the low-dimensional trajectories of M1 (*N_d_* = 4) during the performance of the task. These dimensions explained 54 and 50% of the variance of M1 recordings from monkey C and D, respectively. All networks consisted of N = 50 firing-rate units, similar to previous works [9, 10]. Both types of networks were trained using the same approach followed to train M1 networks. Inputs to SMA networks indicating direction and starting position (*I_dir_*, *I_sp_*) were the same as for M1, except they lasted 500 ms, from 800 to 300 ms before movement onset. The timing of these inputs was designed on the observation of SMA’s dynamics, whose firing rate sharply rises around 800 ms before movement onset. To allow the network to keep track of time as SMA does, we defined a temporal-context input. This input equals 1 from 800 to 300 ms before movement onset and then linearly decreases until the movement ends. The design of the timing of this input was based on the slow decline of the average firing rate of SMA during execution. For the first family of networks (same-control), the weights associated with this input were the same across all conditions and were not changed during training. For the second family of networks (different-control), the temporal-context input entered the network by different sets of weights depending on the movement type (rhythmic or discrete). The sets of weights of discrete and rhythmic movements were designed to be orthogonal to each other and were fixed before training. All networks were trained for 30 thousands epochs. We trained 20 networks of each family for each animal (80 in total). Networks that were not considered successful (1 for same-control, 0 for different-control) were discarded from subsequent analyses.

To test the necessity of the temporal-context input and its structure, extra controls were performed by training RNNs with the same network dynamics (Eqs. 1 and 3) but changing the temporal structure of the temporal-context input (Supp. Fig. 7). To test if this input was necessary, RNNs were trained without temporal-context input, and instead, they received an input to indicate when the movement should stop (Supp. Fig. 7A-C). This set-up is equivalent to the inputs provided to RNNs modelling M1’s dynamics. To test if a decreasing temporal-context input was necessary, we trained RNNs with an input indicating the type of the movement (Supp. Fig. 7D-F) exclusively during preparation. This way, the network is provided with the number of cycles that need to be performed before the movement started. For both analyses, 20 RNNs were trained for each control and animal, giving a total of 80. Supp. Fig. 7 shows two representative solutions found in the trained networks.

RNNs trained to output EMG low-dimensional dynamics (Supp. Fig. 10) used the same inputs described above to specify direction, start position and temporal context. These RNNs were trained using the same training conditions of *N_cycles_* = 4 and 7 cycles for 50,000 epochs.

To simulate a model that keeps the orthogonal inputs for the ends of the rhythmic-to-discrete spectrum but allows continuity for intermediate cases, we rotated the weight vector of the temporal-context input of the already trained Different-control RNNs. The temporal context input of the 7-cycle condition was rotated by an angle (Fig. 4A):

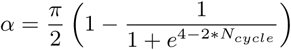

where *N_cycle_* corresponds to the test conditions 1, 2 and 4 cycles. Temporal context inputs for the training conditions (0.5 and 7 cycles) were left intact. RNNs were not retrained to perform the simulations in Fig. 4.

### 4.5 Comparison of RNNs to neural and muscle recordings

We compared two aspects of the RNNs to the neural and muscle recordings. First, to assess the goodness of fit of our models, the outputs of the RNNs and the selected areas (M1 or SMA) were compared for both training and held-out test conditions. For this, we measured how similar the outputs of the RNNs were to the corresponding muscle (Fig. 2B) and neural recordings (Fig. 3B) by using Canonical Correlation Analysis (CCA). CCA outputs a vector of correlations with as many elements as the dimensions of the variables compared. Because the outputs of the RNNs were 4-dimensional, CCs were 4-dimensional, and we summarised these values by their mean. The correlation values range from 0 to 1, with 1 indicating a perfect correlation. CCs were computed with the built-in CCA Matlab function `canoncorr`. To probe how meaningful the CC values were, we estimated the natural similarity between the neural trajectories of the same area as the average canonical correlation between neural trajectories of different animals (grey line in Fig. 2B and 3B).

Second, we used CCA to compare the internal dynamics of the RNNs to the dynamics of the selected area. Low-dimensional trajectories of RNNs (Fig. 2C and Fig. 3C) were extracted by performing PCA across all conditions and applying the same preprocessing as for cortical recordings. Because the RNNs were rate-based networks, neural activity was not smoothed. For these comparisons, only the held-out test conditions were considered (Fig. 2D and 3D). In all CCA comparisons, we considered two 10-dimensional trajectories and summarised the resulting set of correlations by their mean. In addition to estimating the natural similarity of neural trajectories between animals, we estimated the CCs lower bound. We defined the lower bound as the similarity between the neural trajectories of the cortical networks and the neural trajectories of the RNNs reconstructed by projecting RNN activity after first shuffling the order of the 50 dimensions. Neural trajectories from RNNs (timestep Δ*t* = 10 ms) were linearly interpolated to equalise the sampling rate of the neural recordings (Δ*t* = 1 ms).

For both types of comparisons, we tested whether same-control RNNs outperformed different-control RNNs or vice versa. For this, for each test, we performed a two-tailed t-test and a permutation test. Because a t-test determines whether the means of two groups are statistically different, it provides indirect evidence for which family performs better. To provide a direct comparison between the performance of different families, we additionally applied a permutation test that evaluated the percentage of times one family outperformed the other. Here, we compared all possible pairs of CCA values (one element of each RNN family) and divided them by the total number of possible pairs.

### 4.6 Identifying dynamical features of neural trajectories

Neural trajectories of cortical recordings and RNNs were compared in terms of their linear dynamics (Supp. Fig. 2). For each neural trajectory, we selected segments of 360 ms using a sliding window. A linear model for the dynamics ***ẋ*** (*t*) = ***Ax***(*t*) was fitted to each segment, where *x*(*t*) is the N-dimensional vector of trajectory *t* at each time point in the segment and ***A*** is an N x N matrix of parameters. The change in trajectory *ẋ*(*t*) was computed as the first-order difference between adjacent *x*(*t*); we then solved for ***A***. The largest eigenvalue of ***A***, *λ* = *a* + *ib* indicates the most dominant linear dynamics of the trajectory in each segment. If *abs*(*b*) *>* 0 the trajectory rotates. If its real part is negative (*a <* 0), trajectories contract, but if it is instead positive (*a >* 0), trajectories expand. For each population, we used as many dimensions as required to capture at least 50% of the variance. In cases when neural activity required fewer than 3 dimensions, the first 3 dimensions were selected. Since RNNs’ and neural trajectories had different time bases, we divided the eigenvalues of the RNNs by the proportion of the time bases (Δ*t RNN/*Δ*t neural* = 10).

We classified the main dynamical feature as one of 4 possible dynamics according to the values of the real and imaginary parts of the eigenvalues. For this, we used two criteria: First, the absolute value of the imaginary part captures the frequency of the rotations. Thus, considering that the average arm movement had a frequency of *≈*0.5 Hz, we considered that a neural trajectory showed rotational dynamics if its frequency was at least 0.5 Hz (*abs*(*b*) *>* 2 *∗ π/*2). Second, the value of the real part indicates the rate of expansion or contraction. Thus, if neural trajectories expanded/contracted at least 15 % per ms (*a >* 0.0015 and *a < −*0.0015 respectively), we considered the dynamics to be expanding/contracting; and not contracting or expanding otherwise. Then the reported dynamical features correspond to the combination of the characteristics based on the thresholds.

### 4.7 Comparing planes of rotations of neural trajectories

To identify the planes that the neural trajectories rotated on (Supp. Fig. 2B), we applied jPCA [8] by using a publicly available toolbox (at https://churchland.zuckermaninstitute.columbia.edu/content/code). For rhythmic movements, neural activity of 7-cycles movements was selected to define their rotational plane. For each trajectory, only the times corresponding to the pedalling cycles from 2 to 6 were selected to define the plane of rotations. This segment of the neural trajectory corresponds to the steady state [10]. In the case of discrete movements, the rotational plane was defined only using the neural activity during the execution of the half cycle. For each direction and starting position, we computed a pair of planes corresponding to the planes of rotations of neural trajectories of rhythmic and discrete movements. The angle between each pair of planes was computed using the built-in function `subspace` from Matlab.

### 4.8 Distance to the limit cycle

As shown in Fig. 2C and quantified in Supp. Fig. 2, the neural trajectories in M1 during rhythmic movements follow a rotational pattern akin to a limit cycle [9, 10, 12]. To measure how close a given neural trajectory could get to such a limit-cycle (Fig. 2E), we first defined the region it occupied in the subspace. The region the limit-cycle occupied depended on the direction of the movement that was generated. Therefore, to estimate its region, for each direction of a 7-cycle movement, we selected its low-dimensional trajectory from the second to the sixth cycle. The segments corresponding to the neural trajectories of each pedalling cycle were temporally scaled to a fixed length to equalise the number of data points for pedalling cycles. Then, each of the 5 cycles was circularly shifted in time to be aligned to the trajectory of the first selected cycle as closely as possible. The average trajectory of the limit cycle was then defined as the mean of the 5 aligned cycles. The natural variability of the neural trajectories once they reached the limit cycle was defined as a baseline distance (*D_base_*) corresponding to the average distance between the 5 aligned cycles and the average trajectory of the limit cycle.

To measure the Euclidean distance (*D*) between a given trajectory ***X*** of *N_t_* timepoints and *N_D_* dimensions and the limit cycle, the minimum distance between each timepoint of ***X*** and the limit cycle was computed. To compare this distance across RNNs and neural recordings, it was normalised by the natural variability of the limit cycle and the distance to the origin of the subspace as follows:

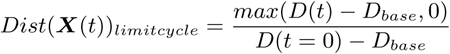

Where *t* = 0 corresponds to 1 s before movement onset (start of movement preparation). This way, the distance to the limit cycle is 1 at the beginning of movement preparation and 0 when the neural trajectory is as close to the limit cycle as the natural variability allows it.

### 4.9 Coding of movement type during movement preparation

We compared how well neural trajectories reflected knowledge of the movement type by analysing how much neural trajectories separated by the upcoming number of cycles during movement preparation (Figs. 2G-H, 3G-H, Supp. Figs. 4 and 8). For this, the Euclidean distance between each pair of neural trajectories was measured for the second leading up to movement onset. We computed this distance between all pairs of trajectories for a set of arm movements that differed only in their number of cycles (black lines), their starting position (grey lines), or their direction (light blue). We then plotted the average distance within each set of pairs in Fig. 2G. To make distances comparable across recordings, all distances were normalised by the average distance between trajectories corresponding to movements of the same number of cycles and directions but opposite starting positions at the time of movement onset. This way, a normalised distance equals 1 when two trajectories are as separated as trajectories of opposite directions. In all analyses, we used the first 6 dimensions of the neural trajectories.

To test if there was a specific subspace that reflected knowledge of the movement type, we performed demixed Principal Component Analysis (dPCA) [15] (Supp. Figs. 5, 9 and 11) across all conditions exclusively on the second leading up to movement onset. dPCA defines the subspaces that best represent chosen task-related variables while maximising the captured variance of the original neural activity. We used the published Matlab package of dPCA (https://github.com/machenslab/dPCA) to perform the analyses. dPCA was applied across all conditions’ averaged neural activity. That is, we used the four marginalisations: Number of cycles, direction, start position and condition-invariant. Neural trajectories of the number of cycles were obtained by projecting the neural activity onto the number-of-cycles axes. The extent of coding of the number-of-cycles was quantified as the sum of the variance explained by all the number-of-cycles axes in the first 15 dPCs of each population.

### 4.10 Initial conditions of movement execution in SMA

To describe the population dynamics of SMA during movement execution, we defined the execution subspace by performing PCA on the neural activity across all conditions and selecting only the times when the movements were being executed (Figs. 3E-F). To estimate the central line of the helix in the low-dimensional subspace, the neural trajectory (*N_d_* = 6) corresponding to 7 cycles was used as a reference (Fig. 3E). For each cycle of the helix, we found its average position in the subspace (grey dots in Fig. 3E). The central line was then defined by fitting a quadratic Bezier curve ***B***(*s*) to the resulting 7 points, where ***B***(*s* = 0) corresponded to the centre of the first cycle (***P***_0_, black dot) and ***B***(*s* = 1) to the centre of the last cycle (***P***_2_, lightest grey dot). This way, the central line is given by

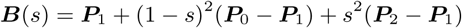

with 0 *≤ s ≤* 1. ***P***_1_ was fitted to minimise the sum of the Euclidean distances between the centres of the helix and the curve ***B***. To estimate the point where each trajectory started along the central line at movement onset, we found its closest point along the central line, ***B***(*s^∗^*). Thus, *s^∗^* corresponded to the position along the central line.

## 5 Data Availability

All data used to produce the figures in this paper (electrophysiological recordings and simultaneous behaviour) are available at [27] DOI: 10.17632/tfcwp8bp5j.1.

## 6 Code Availability

All code used to produce the figures in this paper was developed in MATLAB and Python and is available in the GitHub repository https://github.com/AndreaColinsR/Code_rhythmic_discrete.

## 7 Acknowledgements

ANID Fondecyt Postdoctorado grant 3230117 to A.C.R. and Medical Research Council grant MR/S025944/1 to M.D.H. We thank Abigail Russo and Mark Churchland for making available their data from [9] and Ayeĺen Oyarzo for technical support with code documentation.

## 8 Author Contributions

Conceptualization: A.C.R and M.D.H; methodology: A.C.R. and M.D.H.; investigation: A.C.R.; writing - Original Draft: A.C.R; writing - review and editing: A.C.R, M.D.H and R.F.F.; supervision: R.F.F; resources: A.C.R., M.D.H and R.F.F.;

## Supplementary Information

**Supplementary Figure 1:**
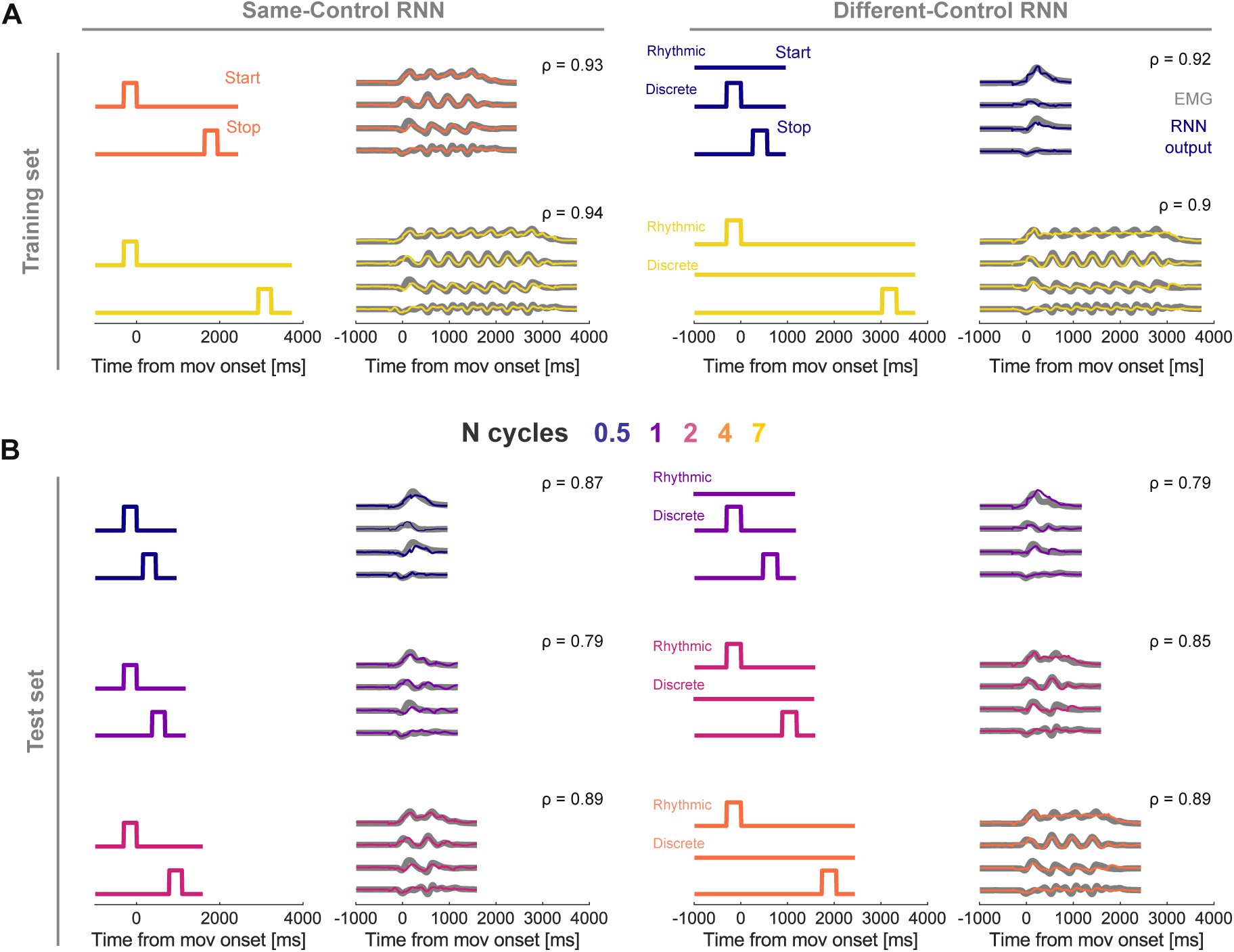
Outputs of trained Recurrent Neural Networks replicate the EMG activity in the train and test datasets. **A)** Left and right panels show the inputs and outputs of an example same-control and different-control network, respectively, for movements in the training dataset. Grey lines show the top four PCs of the EMG that were used to train the networks. Coloured lines are the output of the RNNs. *ρ* values indicate the Pearson correlation between EMG and RNN output for each case. For different-control RNNs, one-hot inputs corresponding to movement type are shown. **B)** As in panel A, but for examples in the test sets of the same RNNs as in panel A

**Supplementary Figure 2:**
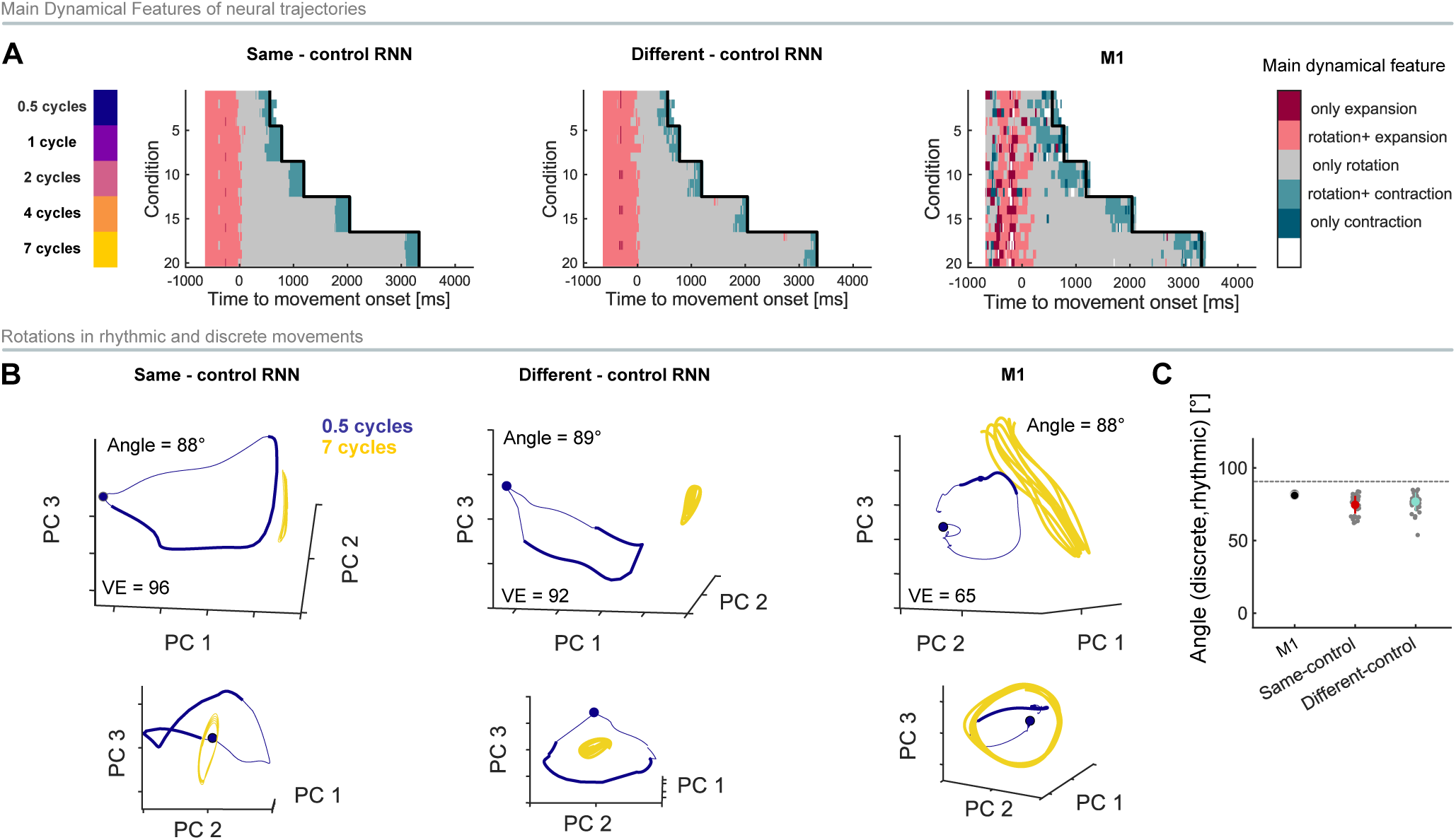
Neural population dynamics in primary motor cortex of rhythmic and discrete movements is underlain by rotations in orthogonal subspaces. **A)** Main dynamical features of the population activity during the arm-pedalling task. Rhythmic and discrete movements are underlain by rotational neural dynamics. Each row corresponds to a given triad of direction, start point and number of cycles. Left colour bar indicates the number of cycles corresponding to each row. Black line shows the movement end. From left to right: Main dynamical features for an example same-control, different-control network and M1 recording. Dynamical features were defined based on the largest eigenvalue of a local linear dynamical system model (see Methods). White regions indicate that a local linear model was not a good fit for those regions (*ρ <* 0.3). **B)** Neural trajectories of discrete (blue) and rhythmic movements (yellow). For the rhythmic movement, only the times of movement execution are shown. For discrete movements, neural trajectories start at 1 s before movement onset and end at 400 ms after the movement ends, and thick lines highlight the times of movement execution. All trajectories correspond to cycling backwards from the cycle’s bottom. Circles show neural state at 1 s before movement onset. Upper and lower panels correspond to different views of the same subspace. VE: variance explained by the top 3 PCs. **C)** Neural trajectories of rhythmic and discrete movements rotate in orthogonal planes. Mean angle between rotational planes of neural trajectories during the execution of discrete and rhythmic movements. Grey dots show the value for each successfully trained network. Error bars show the standard deviation across networks. Grey line indicates the angle at which subspaces are orthogonal.

**Supplementary Figure 3:**
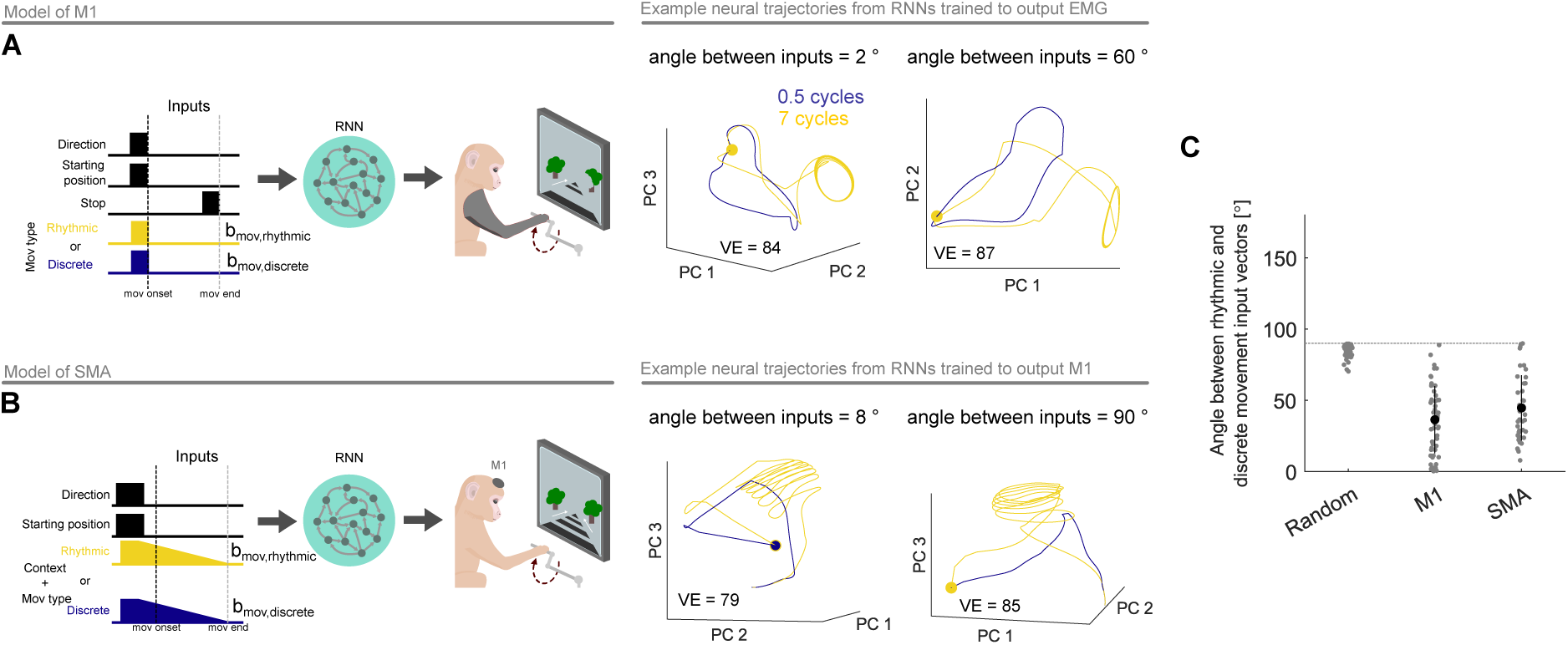
Networks trained to optimise the type of movement input weights show all possible solutions, from same-to different-control. A) Model (left) and example neural trajectories (right) of networks trained to output EMG data. Sets of weights (***b****_mov,rhythmic_*and ***b****_mov,discrete_*) to project movement type inputs were optimised during training. Angle reported is between ***b****_mov,rhythmic_* and ***b****_mov,discrete_*found for the example network. B) As in panel A for SMA model. C) Angle between sets of weights corresponding to discrete (***b****_mov,discrete_*) and rhythmic (***b****_mov,rhythmic_*) movements for all networks trained. Error bars show standard deviation. As a reference, the angle between two random vectors of the same dimensions as the fitted set of weights was measured. Dotted line indicates 90*^◦^*.

**Supplementary Figure 4:**
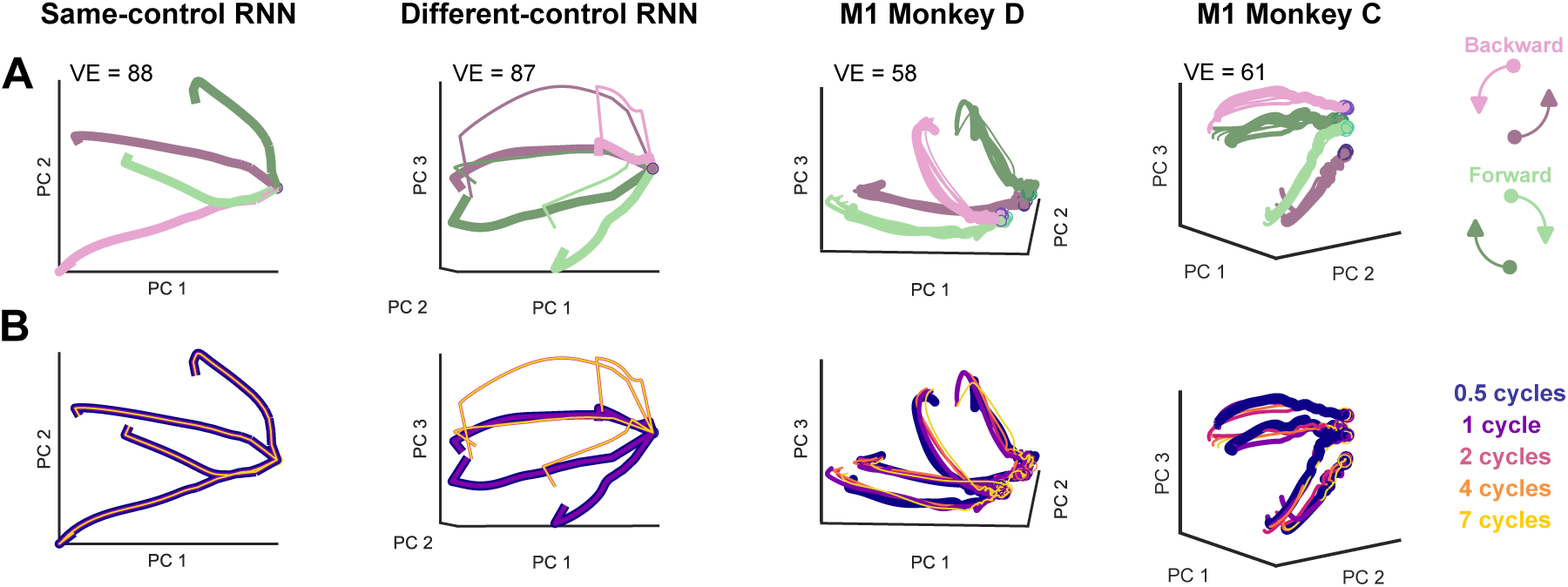
Neural trajectories of M1 and same-control RNNs separate by direction and starting position, but not by number of cycles of the upcoming movement. **A)** Neural trajectories during movement preparation from example same-control RNN, different-control RNN and M1 recordings from both monkeys. In all cases, neural trajectories separated for different directions and starting positions. Trajectories span the interval from 1 s before movement onset up to, but excluding, movement onset. Trajectories corresponding to discrete movements (*N_cycle_* = 1 and 0.5 cycles) are thicker for visualisation purposes only. VE: variance explained by the top 3 PCs. **B)** Same neural trajectories in panel A but coloured by the number of cycles of the upcoming movement. Neural trajectories of different-control RNN were separated by the number of cycles, but not same-control RNNs’ or M1’s. Trajectories corresponding to discrete movements are thicker for visualisation purposes.

**Supplementary Figure 5:**
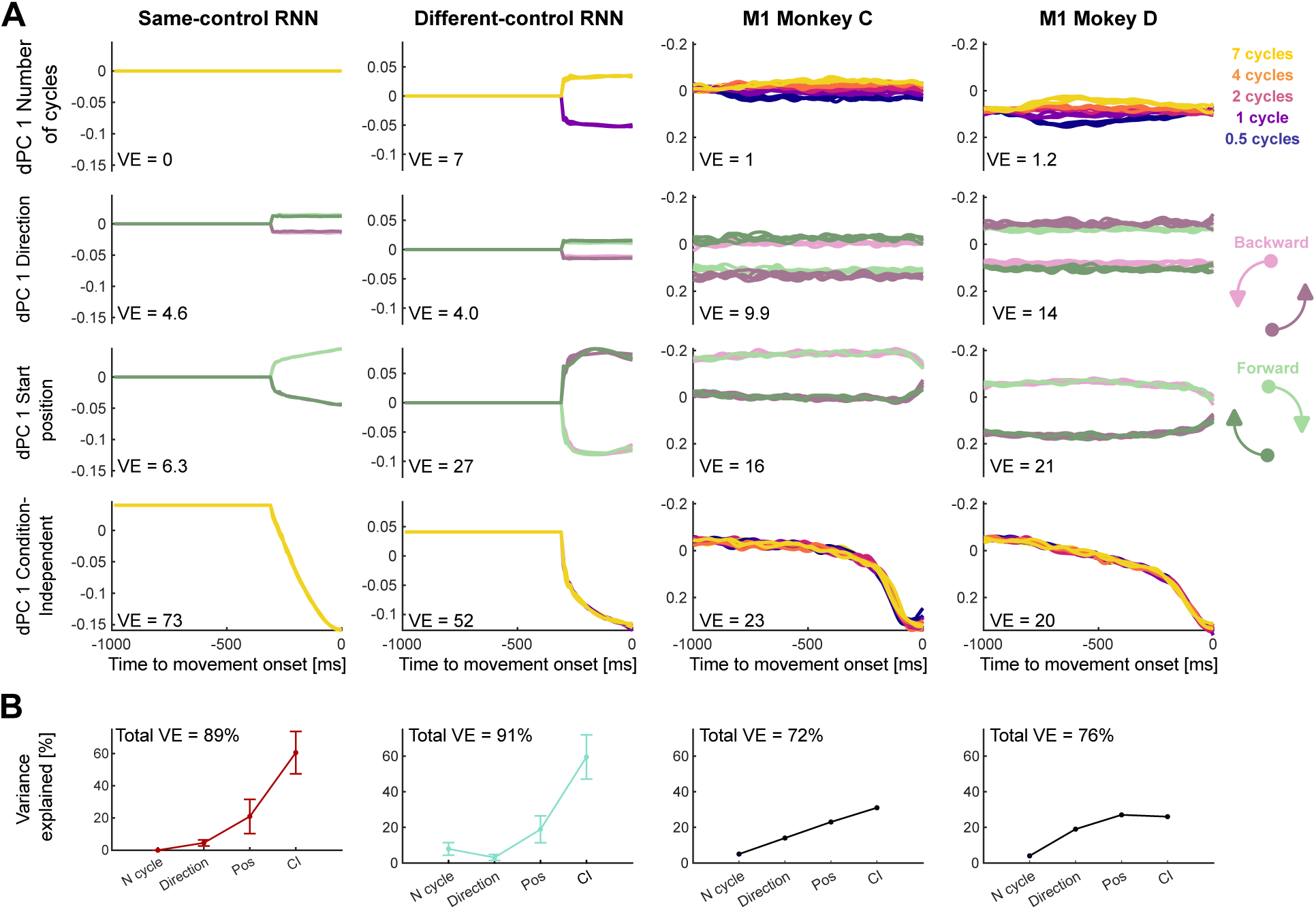
The number of cycles of the upcoming movement is the least encoded variable in M1 and Same-control RNNs. **A)** Decoding axes for each movement feature determined by dPCA during movement preparation. From top to bottom: number-of-cycles-, direction-, start position- and Condition-Invariant dPCs. From left to right: Trajectories from example same-control RNN, different-control RNN and trajectories from M1 recordings from both animals. Variance Explained in percentage (VE) is shown for each dPC. The range of dPCs values is comparable across rows. **B)** Variance Explained by the subspaces coding for number of cycles, direction, start position and Condition-Invariant. The total variance explained was calculated as the sum of the variance explained in the first 15 dPCs. Because RNNs are noiseless, these dPCs account for nearly all variance in their activity. By contrast, the variance in M1’s activity includes contributions from sources other than the tested conditions, so a lower amount of variance can be captured by these dPCs. Error bars show the SD across all successfully trained RNNs.

**Supplementary Figure 6:**
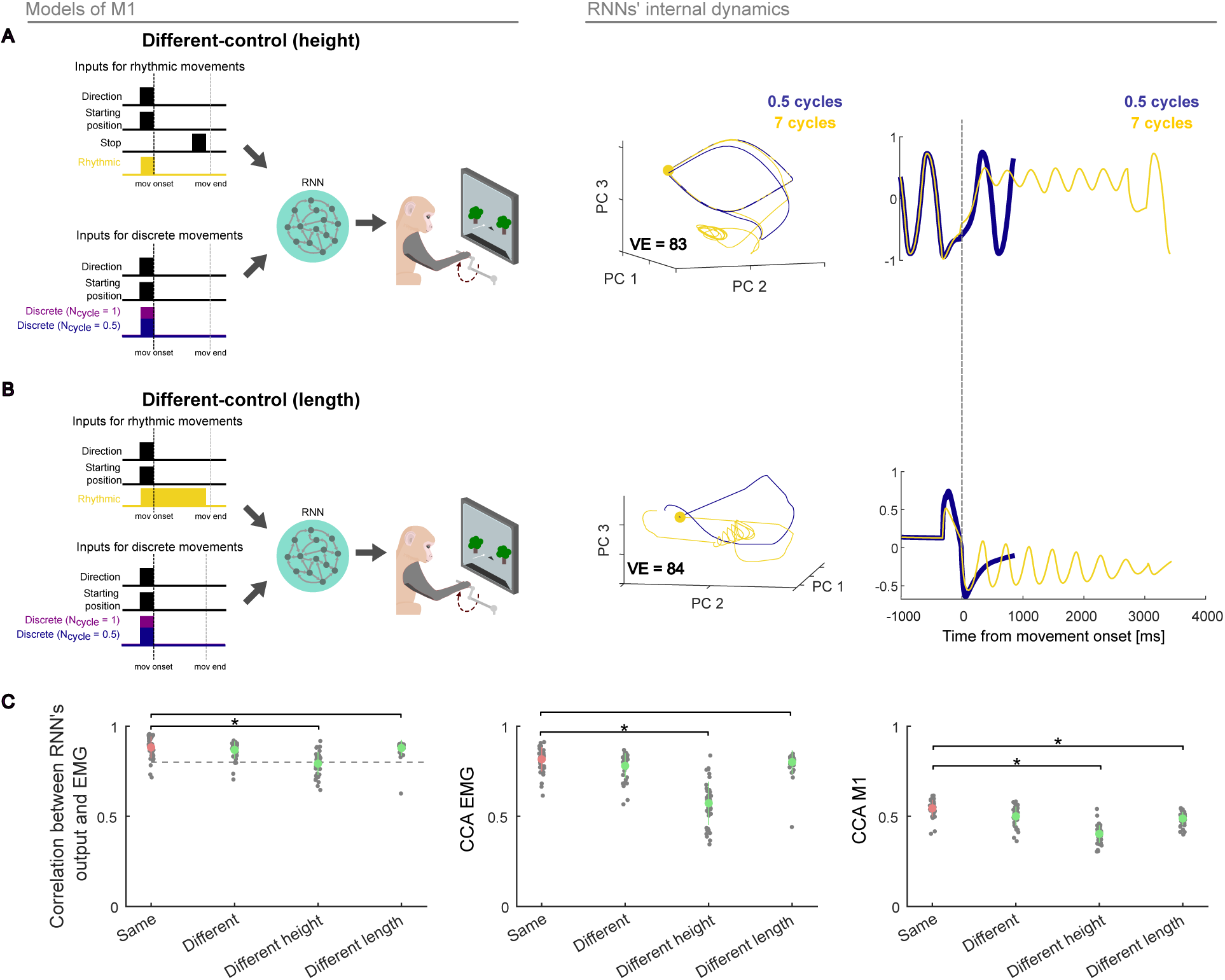
Variants of different-control models trained to output EMG trajectories show dynamics incompatible with M1’s. Two variants of the different-control (**A)** and **B)**) were trained to output the low-dimensional activity of EMG. **A)** Left: RNNs received a one-hot-coded input indicating if movement was rhythmic or discrete. Rhythmic movements received a stop signal, but discrete movements did not. Instead, for discrete movements, the number of cycles was encoded in the height of the discrete-movement input. Central and right panels show that an RNN of this type had internal dynamics incompatible with M1: before movement onset, the RNN’s trajectories formed a limit cycle whereas M1’s trajectories did not. **B)** Same as panel A for the other variant control model: For rhythmic movements, the RNN received a constant input until 100 ms before the end of the movement, while discrete movements received an input only during preparation. For discrete movements, the number of cycles is encoded in the height of this input. The internal dynamics of these RNNs showed a rotational trajectory of decreasing amplitude for rhythmic movements, inconsistent with M1 dynamics. **C)** From left to right: Pearson correlation between EMG and RNNs’ outputs, CCA between EMG and RNNs’ outputs, and CCA between M1 dynamics and internal dynamics of RNNs. Dashed horizontal line shows the threshold used to classify the networks as successful. Same and Different models correspond to same- and different-control models in Figure 2. RNNs from the variant of different-control models were significantly less similar to M1 than the original same-control model (right). * denotes p-value*<* 0.05, t-test (N = 40). In our permutation test, the same-control model outperformed different-height (EMG = 96%, M1 = 96%) and different-length models in most cases (EMG = 60%, M1 = 86%).

**Supplementary Figure 7:**
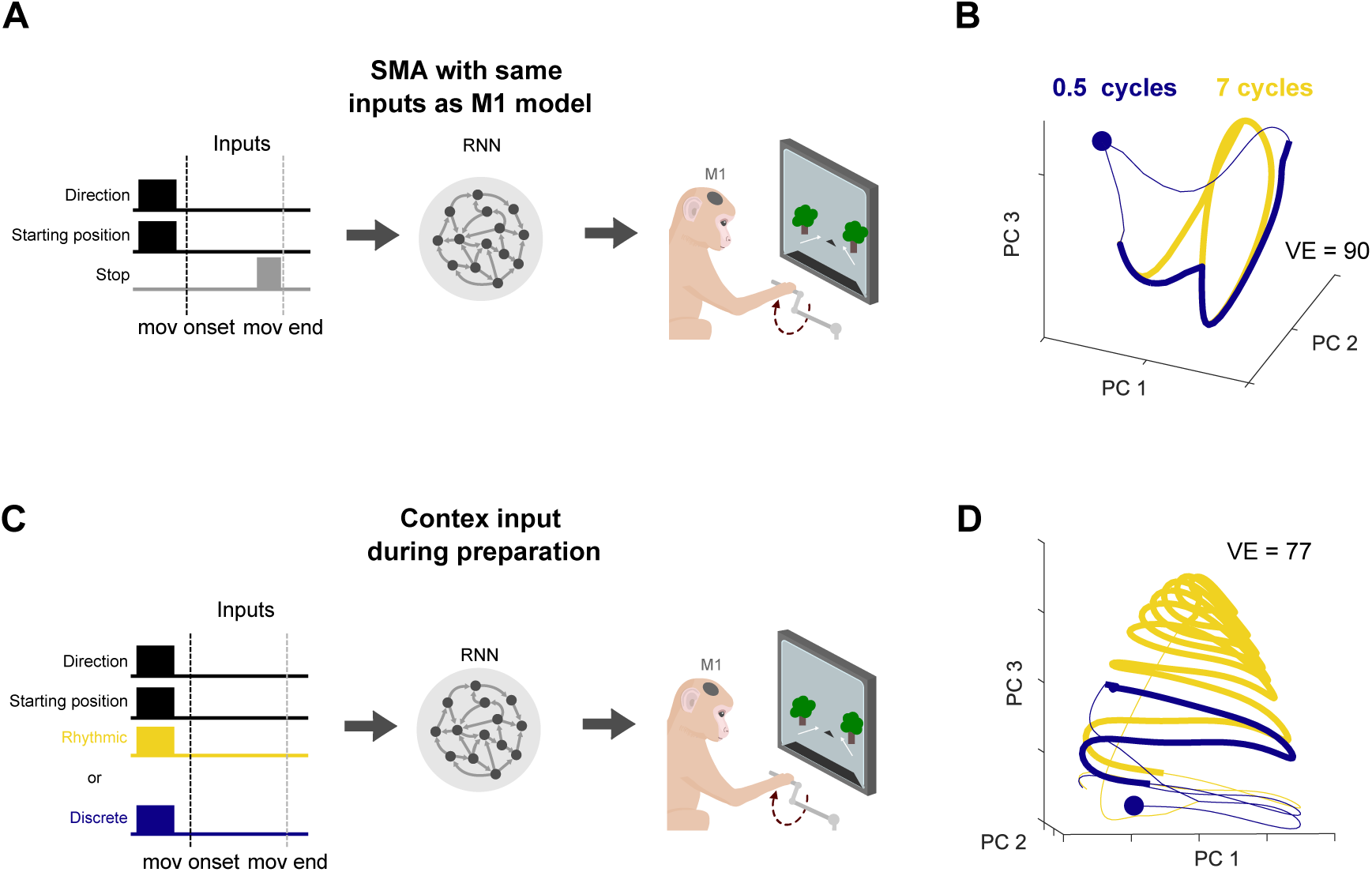
Temporal-context input is key to model the population dynamics of SMA. Models without or with a variant of the temporal-context input used in RNNs Fig. 3 solved the task but did not capture the characteristic helical structure in SMA dynamics. **A)** Schematic of the trained RNNs without a temporal-context input. Inputs for these networks were the same as the inputs used to model the dynamics of M1. **B)** Neural trajectories from an example RNN trained with inputs defined in A. Thick lines highlight times during movement execution. The dynamics of RNNs lacking the temporal-context input (like networks in Fig. 2A) were governed by a limit-cycle. **C)** Schematic of the trained RNNs receiving the number of cycles only during movement preparation. Inputs from discrete and rhythmic movements entered the network through orthogonal sets of weights. In contrast to the models in Fig. 3A, the networks did not receive decreasing inputs during the movement execution. **D)** Neural trajectories from an example RNN trained with inputs defined in C. Thick lines highlight times during movement execution. Neural trajectories show a spiral instead of the helical structure as expected for SMA.

**Supplementary Figure 8:**
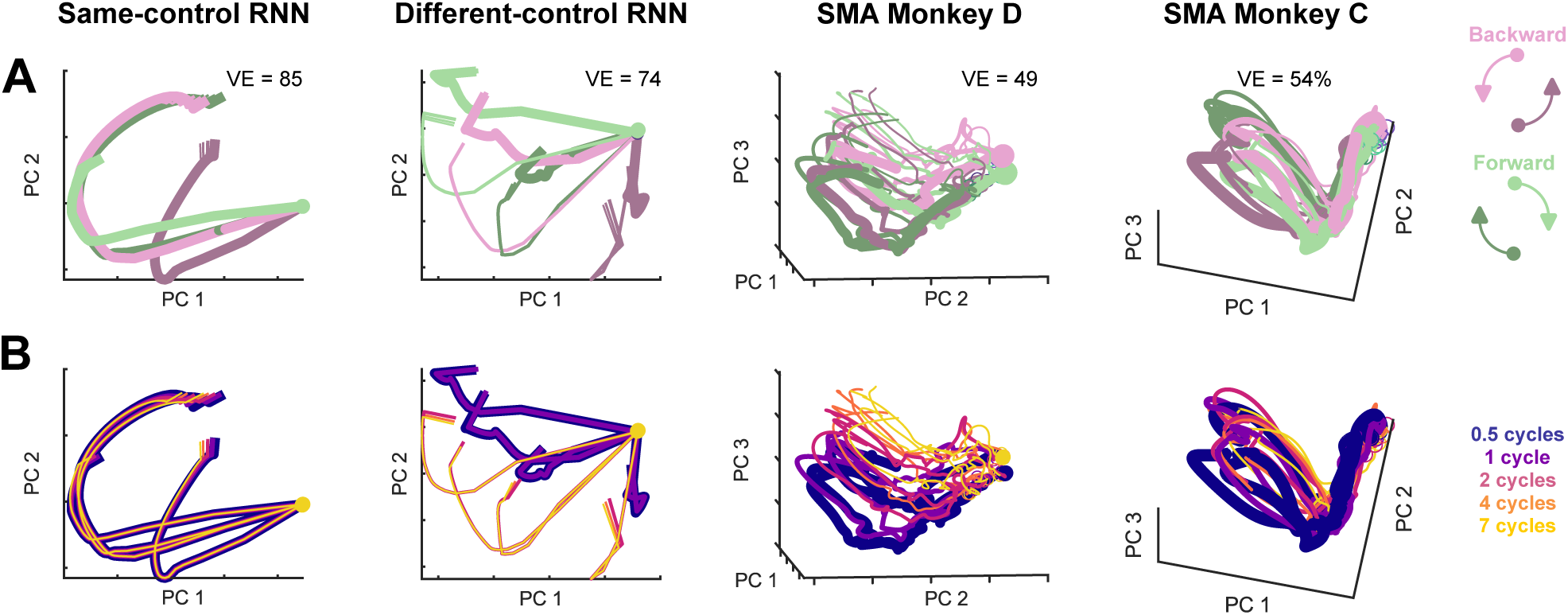
Neural trajectories of SMA and different-control RNNs separated by the number of cycles of upcoming movement. **A)** Neural trajectories during movement preparation from example same-control RNN, different-control RNN and SMA recordings from both monkeys. VE: variance explained by the top 3 PCs. **B)** Same neural trajectories in panel A, but coloured by the number of cycles of the upcoming movement.

**Supplementary Figure 9:**
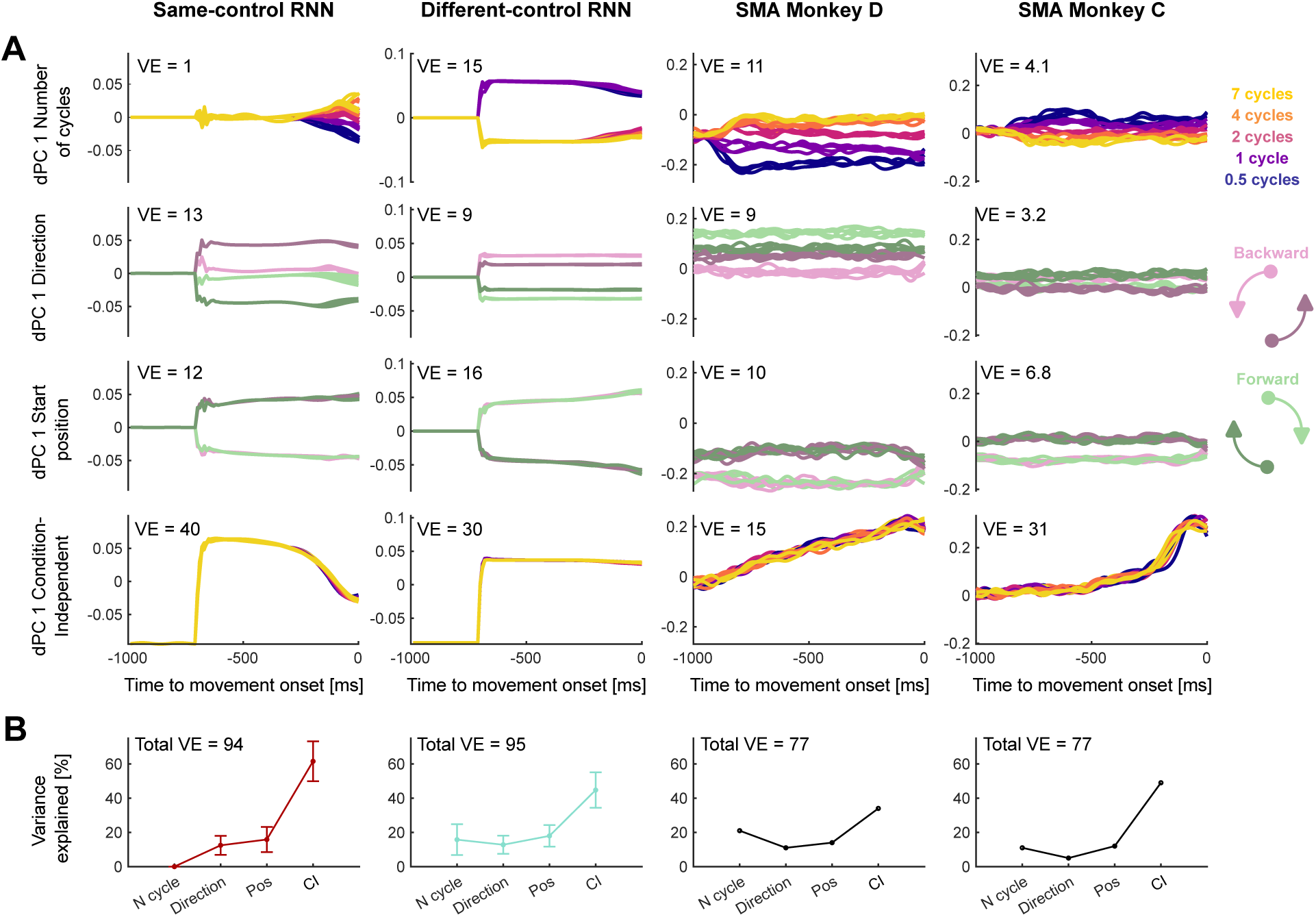
The number of cycles of the upcoming movement is encoded in SMA and different-control RNNs. **A)** From top to bottom rows: First number-of-cycles-, direction-, start position- and Condition-Invariant dPCs of an example Same-control (left) RNN, a Different-control (middle) RNN and SMA (right) during movement preparation. Variance Explained (VE) in percentage is shown for each dPC. The range of dPCs values is comparable across rows. **B)** Variance Explained by the subspaces coding for the number of cycles, direction, start position and Condition-Invariant. The total variance explained was calculated as the sum of the variance explained in the first 15 dPCs.

**Supplementary Figure 10:**
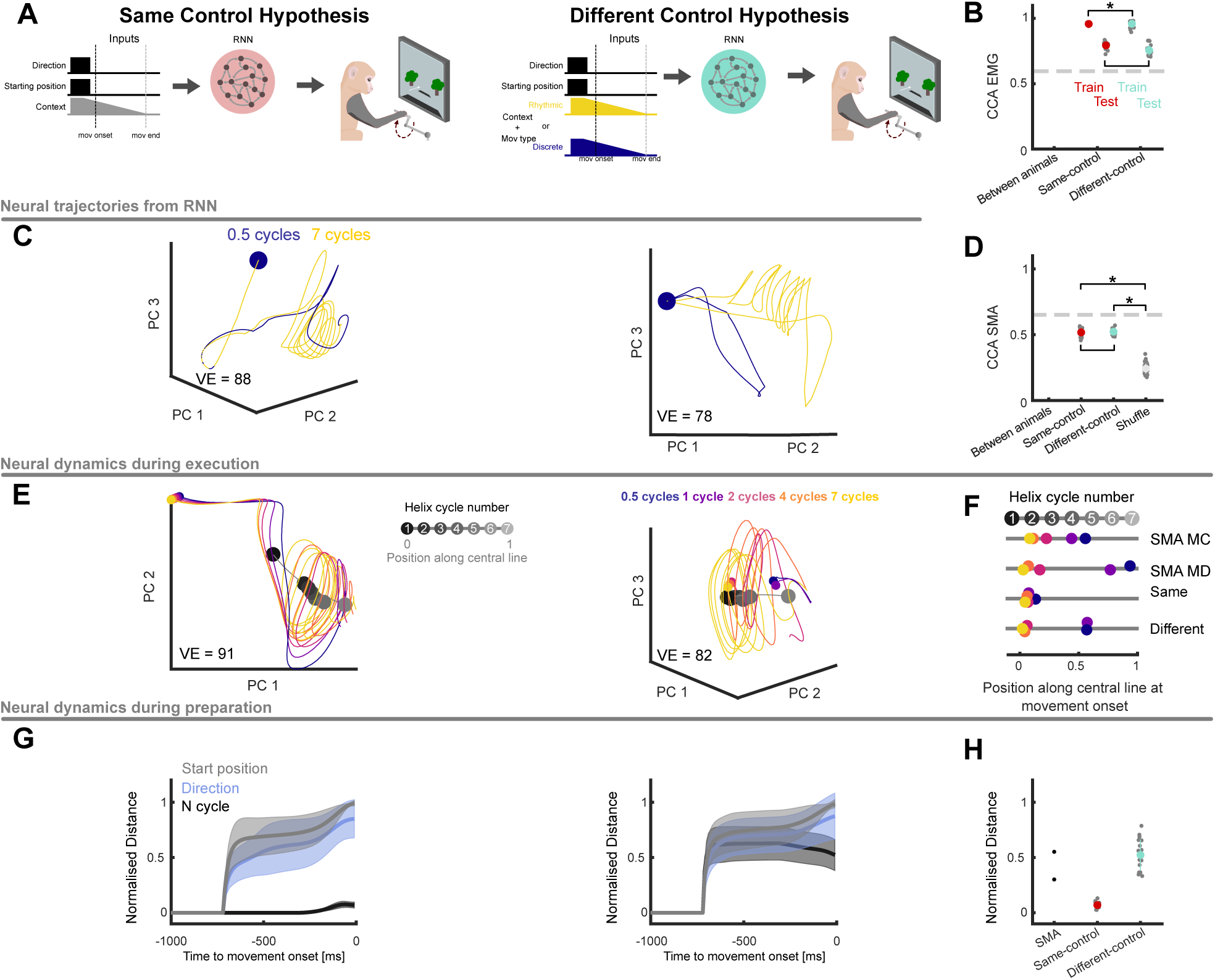
Neural networks trained to output EMG trajectories using SMA inputs show the same properties of SMA. **A)** Two families of RNNs were trained to output the low-dimensional activity of EMG using the same inputs as SMA models. Left: same control strategies for all movements. Right: Different control strategies for distinct movements. RNNs received the same inputs as the same-control RNNs, but the temporal-context input specified the type of movement (discrete or rhythmic). **B)** Similarity (CCs) between EMG trajectories and RNNs’ output. Each grey dot corresponds to one RNN. Error bars show the standard deviation across RNNs. * denotes a significant difference between two quantities (t-test, p-value*<*0.05). **C)** Example neural trajectories from discrete and rhythmic movements. All trajectories correspond to pedalling forward from the bottom. Circles show the start of movement preparation. **D)** Similarity (CCs) between SMA trajectories and internal dynamics of RNNs. Comparisons were made for SMA recordings across animals, SMA versus the dynamics of the trained RNNs and SMA versus the dynamics of trained RNNs whose projections into neural dimensions were shuffled. * denotes a significant difference between two quantities (t-test, p-value*<*0.05). Horizontal bar without * indicate no significant difference. **E)** Example neural trajectories of a same- and different-control RNN (left and right, respectively) during the execution of rhythmic and discrete movements. Coloured circles indicate the trajectories’ positions at movement onset for different numbers of cycles. Black to grey circles show the average activity for each cycle of the 7-cycle movement. Grey line shows the central line of the helix, which was fitted to the centres of the cycles. **F)** Mean position along the central line of the helix of the neural trajectories at the onset of different cycling movements. Vertical position of each dot was randomised around the grey line for visualisation purposes only. **G)** Distance during preparation between neural trajectories corresponding to different movement features. All distances were normalised by the distance between the neural trajectories at the onset of movements that started in opposite positions (same direction and number of cycles). Solid lines, mean across RNNs or conditions (SMA); shaded area, standard deviation across RNNs. **H)** Normalised distance between trajectories corresponding to different number of cycles at movement onset.

**Supplementary Figure 11:**
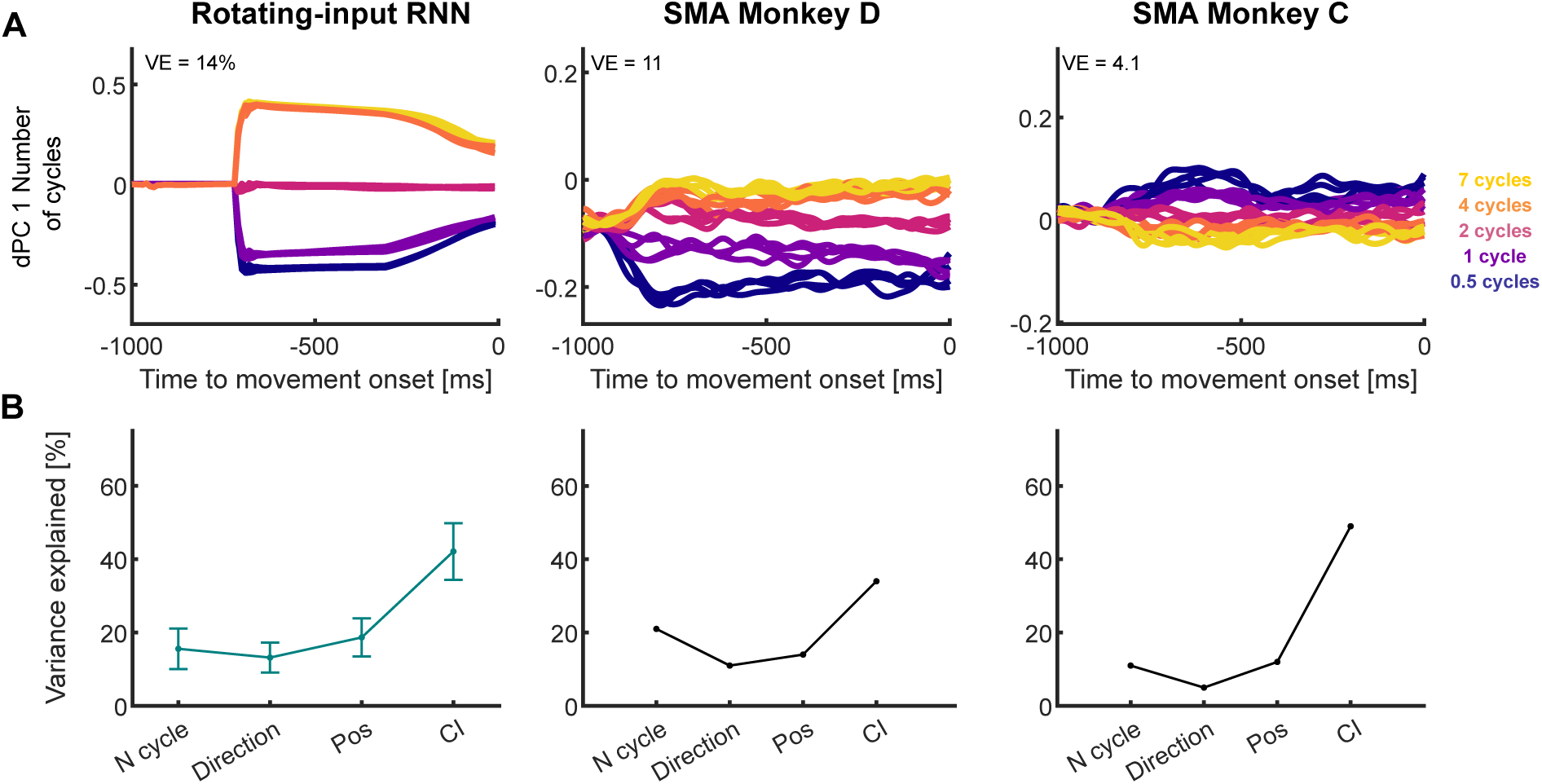
Rotating-input RNN captures sorting of neural trajectories by number of upcoming cycles. **A)** Activity projected onto the dominant number-of-cycles axis found by dPCA, for rotating-input RNN (left) and SMA activity (centre and right; redrawn from Supp. Fig. 9). **B)** Variance explained by dPCA-obtained subspaces of preparation activity coding for the number of upcoming cycles, direction, start position, and the Condition-Invariant (CI) subspace, for rotating-input RNNs (left) and SMA (centre and right; redrawn from Supp. Fig. 9).

Supplementary Video 1: Neural trajectories from both RNN families and M1. For Same- and Different-control RNNs: black line at the bottom shows the timing of the direction input, blue and yellow lines show the timing of the stop inputs for discrete and rhythmic movements, respectively.

Supplementary Video 2: Neural trajectories from both RNN families and SMA. For Same- and Different-control RNNs: black line at the bottom shows the timing of the direction input, blue and yellow lines show the timing of the temporal context inputs for discrete and rhythmic movements, respectively.

